# MISPEL: A supervised deep learning harmonization method for multi-scanner neuroimaging data

**DOI:** 10.1101/2022.07.27.501786

**Authors:** Mahbaneh Eshaghzadeh Torbati, Davneet S. Minhas, Charles M. Laymon, Pauline Maillard, James D. Wilson, Chang-Le Chen, Ciprian M. Crainiceanu, Charles S. DeCarli, Seong Jae Hwang, Dana L. Tudorascu

## Abstract

Large-scale data obtained from aggregation of already collected multi-site neuroimaging datasets has brought benefits such as higher statistical power, reliability, and robustness to the studies. Despite these promises from growth in sample size, substantial technical variability stemming from differences in scanner specifications exists in the aggregated data and could inadvertently bias any downstream analyses on it. Such a challenge calls for data normalization and/or harmonization frameworks, in addition to comprehensive criteria to estimate the scanner-related variability and evaluate the harmonization frameworks. In this study, we propose MISPEL (Multi-scanner Image harmonization via Structure Preserving Embedding Learning), a supervised multi-scanner harmonization method that is naturally extendable to more than two scanners. We also designed a set of criteria to investigate the scanner-related technical variability and evaluate the harmonization techniques. As an essential requirement of our criteria, we introduced a multi-scanner matched dataset of 3T T1 images across four scanners, which, to the best of our knowledge is one of the few datasets of this kind. We also investigated our evaluations using two popular segmentation frameworks: FSL and segmentation in statistical parametric mapping (SPM). Lastly, we compared MISPEL to popular methods of normalization and harmonization, namely White Stripe, RAVEL, and CALAMITI. MISPEL outperformed these methods and is promising for many other neuroimaging modalities.

## 1 Introduction

There is a growing interest in the neuroimaging community in combining imaging data from a variety of diverse datasets to enable large-scale multi-study analyses that have high statistical power, reliability, and robustness (Madan, 2021; Mar et al., 2013; Madan, 2017; Milham et al., 2018). Despite the promise of massive data aggregation initiatives, large-scale neuroimaging analyses from such data collections often suffer from issues of technical variability due to scanner- and individual-based heterogeneity across studies, which may introduce bias in imaging-derived measures (Kruggel et al., 2010; Potvin et al., 2019; Torbati et al., 2021a) and causes alterations of the biological signals of clinical interest (Shinohara et al., 2014a, 2017), among other unwanted and unexpected artifacts. Scanner technical variability has been majorly recognized as intensity unit effects and scanner effects (Wrobel et al., 2020; Torbati et al., 2021a).

Intensity unit effects are due to the arbitrary nature of image intensity scale, which can cause variability in interpretations of intensity units and thus make the direct quantitative analysis of image intensities difficult (Wrobel et al., 2020). Intensity unit effects have been long recognized and addressed by intensity normalization methods, such as White Stripe (WS) (Shinohara et al., 2014b), a well-known normalization method in neuroimaging. A comprehensive review of the initial intensity normalization methods can be also found in (Shah et al., 2011).

Scanner effects refer to any post-normalization inter or intra-scan variation that is not biological in nature (Fortin et al., 2016) and stems from scanner and acquisition differences (Dinsdale et al., 2021). So far, these causes of variation have been recognized: differences in scanner manufacturer (Takao et al., 2014), scanner upgrade (Han et al., 2006), scanner drift (Takao et al., 2011), scanner field strength (Han et al., 2006), and gradient non-linearities (Jovicich et al., 2006). An example of such effects can be seen in tissue type volumes extracted from White Stripe (WS)-normalized images in Figure 1b. The group of methods that aim to remove scanner effects is referred to as harmonization. Harmonization is a complex and challenging task due to (1) lack of thorough understanding of scanner effects, and (2) lack of standardized criteria for assessment of scanner effects and evaluation of harmonization.

**Figure 1:**
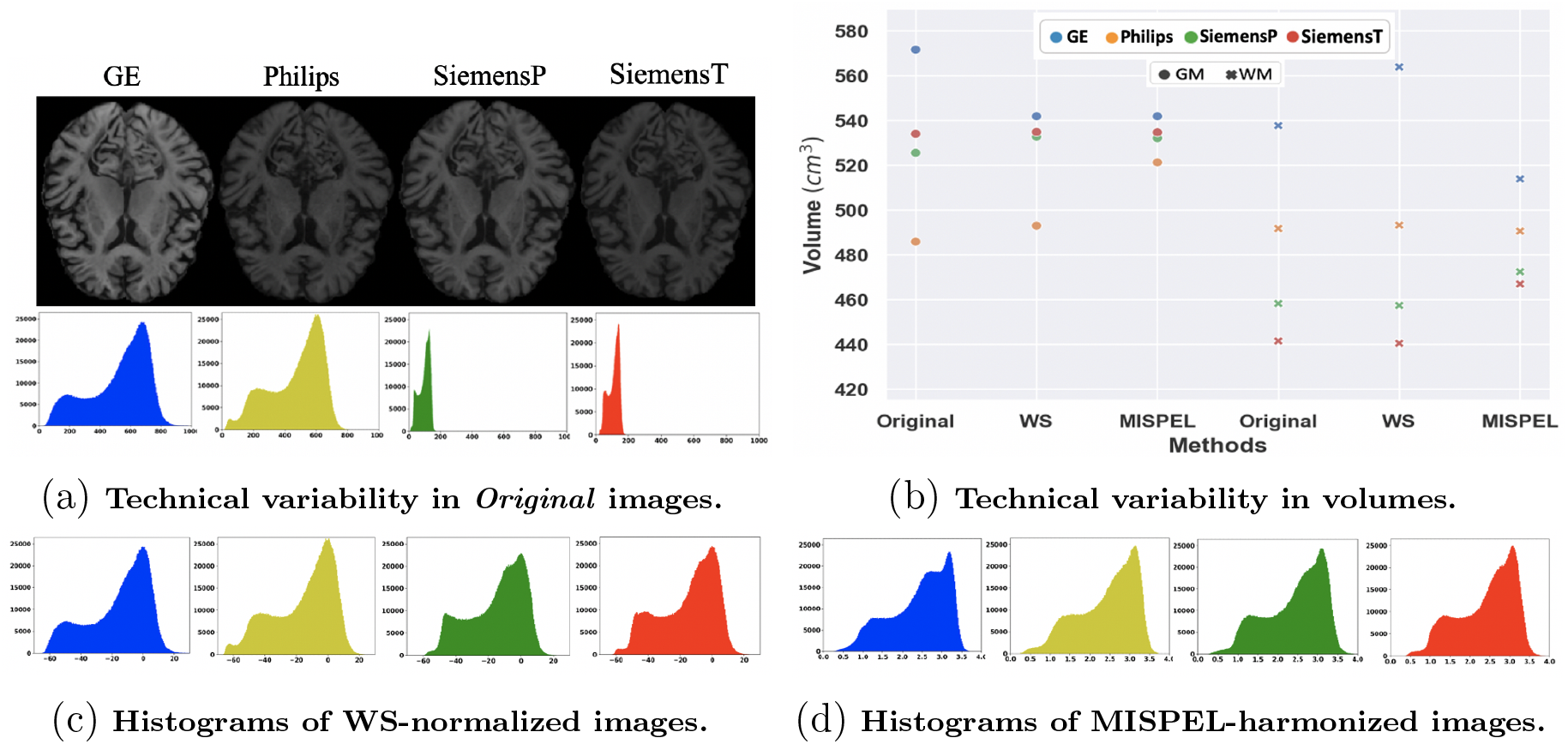
Example of technical variability, White Stripe normalization, and MISPEL harmonization in *matched* images. For this example, *Original* images are the matched T1 MRIs scanned on four different 3T scanners: GE, Philips, SiemensP, and SiemensT. Specifications of these scanners can be found in Table 1. Figure (a) depicts the technical variability of the Original images as dissimilarity in contrast of their axial slices, and discrepancy among histograms of their whole brain. Figure (b) shows the technical variability of matched images in terms of their tissue type volumetric dissimilarity. The volumes were depicted for the Original images as well as their WS-normalized and MISPEL-harmonized versions. Figures (c) and (d) depict the histograms of whole brain in WS-normalized and MISPEL-harmonized matched images, respectively. Histograms of matched images have identical axes and correspond (from left to right) to GE, Philips, SiemensP, and SiemensT scanners. For interpretation of the references to color in this figure legend, the reader is referred to the web version of this article.

In this specific study, our main interest lies in understanding and compensating for technical variability of images, specifically the scanner effects. Scanner effects cannot be easily removed by simple intensity distribution matching (Fortin et al., 2016) or a linear transformation of images (Wrobel et al., 2020). Even though there has been a noticeable growth in the number of studies focused on scanner effects and harmonization recently (Dewey et al., 2019, 2020; Liu et al., 2021; Cackowski et al., 2021; Zuo et al., 2021), there is a lack of insight with respect to how these scanner effects appear on images. One main reason could be the lack of ground truth for these studies, which leaves them with no standard evaluation criteria and consequently makes their observations partly incoherent and hard to compare. Based on the observations confirmed by several of these studies, it is now known that scanner effects can vary across the voxels of an individual image (Chen et al., 2020a). Furthermore, it is also known that scanner effects change the tissue contrast and consequently affect the results of tissue segmentations (Meyer et al., 2019). Torbati et al. (2021a) has shown that scanner effects can affect different regions of brain differently and result in regional summary measures with varying degree of scanner effects.

The best experimental design setup to understand and quantify scanner effects is to conduct a paired study by having subjects travel to different sites/scanners, to collect the *paired* dataset (Dewey et al., 2019; Zuo et al., 2021). A paired dataset is a set of *paired* images that are the images of each individual scanned on *two* scanners with short time gap. Paired images are expected to be images of biologically similar brain with differences solely due to scanner effects. Using a paired dataset, scanner effects and harmonization can be estimated as similarities and dissimilarities within paired images, respectively. As such, a ground truth is not necessary.

Figure 1 illustrates an example from a *matched* dataset, similar to a paired dataset but with more than two scanners. An example of technical variability across MRI scanners can be observed as dissimilar contrast and voxel intensity histograms of these matched images in Figure 1a, as well as the resulting variability in deduced volumes for both gray matter (GM) and white matter (WM) tissue types in Figure 1b. Also, Figure 1c depicts the histograms of the WS-normalized version of the matched images. The scanner effects can be observed in the WS-normalized images as their dissimilar histograms in Figure 1c, as well as their discrepant volumes in Figure 1b.

From a methodological and more specifically, a machine learning perspective, paired and unpaired datasets are considered respectively as labeled and unlabeled data for the task of harmonization. Accordingly, the harmonization methods developed based on paired and unpaired data are called the supervised and unsupervised methods (Dewey et al., 2019; Zuo et al., 2021; Torbati et al., 2021a; Liu et al., 2021). The majority of the research on harmonization is currently focused on the unsupervised methods, due to scarcity of matched or even paired datasets. However, there exist two supervised methods, namely DeepHarmony (Dewey et al., 2019) and mica (Wrobel et al., 2020). DeepHarmony is a contrast harmonization method that maps images of two scanners to a middle-ground space in which images are harmonized by having similar contrast. However, DeepHarmony is limited to harmonizing images of just two scanners. On the other hand, mica is a multi-scanner (i.e., more than two scanners) method that harmonizes images by adapting their intensity distributions to that of the *target* scanner. Even though adapting images to a target scanner seems to simplify the task of harmonization, it introduces the new challenge of determining the “best” scanner in the pooled data. Selecting such scanner is not a trivial task when, for example, motion artifacts in images could be of concern (Alexander-Bloch et al., 2016; Torbati et al., 2021a).

Harmonization can be applied to, two broad categories: (1) harmonization of image-derived measures, and (2) harmonization of images. The methods of the first category can be described as ComBat (Johnson et al., 2007) and its extensions (Beer et al., 2020; Chen et al., 2020b; Pomponio et al., 2020; Reynolds et al., 2022). ComBat is a location and scale adjustment method used in neuroimaging for harmonizing image-derived measures and has been applied to images of different modalities: DTI (Fortin et al., 2017), MRI (Fortin et al., 2018), and fMRI (Nielson et al., 2018). Even though ComBat is a straightforward method which showed success in harmonization of image-derived measures in many studies (Yu et al., 2018; Radua et al., 2020; Foy et al., 2020; Torbati et al., 2021a), its performance cannot be easily evaluated at the image level. Moreover, ComBat is directly affected by the known or unknown biological differences among subjects. In fact, ComBat is prone to removing the biological variability that is correlated to scanner effects and was not known to be considered through ComBat harmonization (Liu and Markatou, 2016; Obenauer et al., 2019). Thus, a potentially better way to approach harmonization is to conduct it at the image level.

RAVEL was proposed as the first normalization and harmonization framework, removing inter-subject variability from MRIs at the voxel level (Fortin et al., 2016). Harmonization methods using deep learning techniques have subsequently been proposed. The unsupervised deep-learning-based methods treat harmonization as the task of domain or style transfer learning, in which images of scanners are mapped to the domain or style of one selected scanner, called *target* scanner (Dewey et al., 2020; Zuo et al., 2021). As well as the challenge of selecting the target scanner, these methods have other limitations based on the deep learning network they used for transfer. For example, methods using CycleGAN (Modanwal et al., 2020) or DualGAN (Zhong et al., 2020) networks are limited to harmonization of just two scanners. Another example is CALAMITI with a disentanglement network limited to harmonizing inter-modality paired dataset. These data consist of paired images of two predetermined modalities taken from an individual on the *same* scanner with a short time gap. Methods using style transfer (Liu et al., 2021; Liu and Yap, 2021) are prone to mapping images to the biological or clinical information of the target scanner if images across scanners are confounded by this information. Another major group of unsupervised methods proposed generating scanner-invariant latent representations for synthesizing harmonized images (Moyer et al., 2020) or training the neuroimaging tasks on images (Aslani et al., 2020; Dinsdale et al., 2021). However, these methods are prone to lose the information of images during harmonization, as their generated representation has been proven to be limited to the least informative scanner (Moyer and Golland, 2021).

In this work, we present MISPEL (**M**ulti-scanner **I**mage harmonization via **S**tructure **P**reserving **E**mbedding **L**earning), which is a supervised multi-scanner harmonization method that maps images of scanners to a middle-ground harmonized image space.

Figures 1d and b depict the result of MISPEL on harmonizing our example of matched images. In this study, we also introduce a multi-scanner matched dataset of 3T T1 images across four scanners, one of the few datasets of its kind (Duchesne et al., 2019; Magnotta et al., 2020; Maikusa et al., 2021; Hawco et al., 2022). In addition, we provide a set of experiments assessing scanner effects and evaluating harmonization on our unique set of matched data by applying commonly used MR image processing and segmentation software packages FSL (Zhang et al., 2001), SPM (Ashburner and Friston, 2005), and FreeSurfer (Fischl, 2012). Lastly, we compare MISPEL with three well-known methods of image-based normalization and harmonization, White Stripe, RAVEL, and CALAMITI.

## 2 Materials and methods

### 2.1 Study population and image acquisition

The sample used in this study consists of 18 participants which are part of an ongoing project (UH3 NS100608 grant to J. Kramer and C. DeCarli). The median age of the participants was 72 years (range 51-78 years) and 44% (N = 8) were males. All participants were cognitively unimpaired with either a low or high degree of small vessel disease (SVD) as previously defined (Wilcock et al., 2021)). T1-weighted (T1w) images were acquired for each participant on each of four different 3T scanners [GE, Philips, SiemensP, and SiemensT (Table 1)]. For each participant, these matched images were taken at most four months apart, a time period over which we assume no biological changes could occur in the brain and differences observed between any pairs of scans are solely due to the scanner effects. In a matched dataset, the scanner and harmonization effects can be estimated based on the dissimilarity and similarity of matched images, respectively. The details of estimation of scanner effects and evaluation of harmonization methods are provided in Section 2.5.

**Table 1:** Scanner specifications.

| Scanner Name | GE | Philips | SiemensP | SiemensT |
| --- | --- | --- | --- | --- |
| Manufacturer | General Electrics | Philips | Siemens | Siemens |
| Scanner Hardware | DISCOVERY-MR750w 3T | Achieva-dStream 3T | Prisma-fit 3T | TrioTim 3T |
| Scanner software | 27-LX-MR-Software-release:<br>DV26.0-R03-1831.b | 5.6.1-5.6.1.0 | syngo-MR-E11 | syngo-MR-B17 |
| Receive Coil | 32Ch-Head | MULTI-COIL | BC | 32Ch-Head |
| T1-w Sequence Type | BRAVO | ME-MPRAGE | ME-MPRAGE | ME-MPRAGE |
| Resolution (mm) | $1.0 \times 1.0 \times 0.5$ | $1.0 \times 1.0 \times 1.0$ | $1.0 \times 1.0 \times 1.0$ | $1.0 \times 1.0 \times 1.0$ |
| TE/ΔTE (ms) | 3.7 | 1.66/1.9 | 1.64/1.86 | 1.64/1.86 |
| TR (ms) | 9500 | 2530 | 2530 | 2530 |
| TI (ms) | 600 | 1300 | 1100 | 1200 |

### 2.2 Image preprocessing

We use RAVEL as one of our harmonization methods in this study. In order to prevent confounding our evaluation with inconsistent preprocessing steps, we preprocessed all images using the pipeline prescribed for RAVEL (Fortin et al., 2016). Therefore, we first used a non-linear symmetric diffeomorphic image registration algorithm (Avants et al., 2008) to register images to a high-resolution T1-w image atlas (Oishi et al., 2009). We then applied the N4 bias correction method (Tustison et al., 2010) to the registered images to correct them for spatial intensity inhomogeneity. As the last step of the pipeline, we skull stripped the images using the mask provided in (Fortin et al., 2016). We also scaled images in one additional step, in which intensity values of each image were divided by their within-mask average intensity value. Throughout this manuscript, these preprocessed images are referred to as *RAW* and used as input to our models.

### 2.3 MISPEL

Our proposed framework, MISPEL, is a convolutional deep neural network for harmonizing images from multiple scanners, for which a *matched* dataset is available. We designed MISPEL to (1) generalize to multiple (more than two) scanners, (2) preserve the structural (anatomical) information of the original brains, (3) learn harmonization on a matched dataset, and (4) later harmonize unmatched images of the scanners for which the matched dataset was collected. Although it is more desirable to train a harmonization method on the whole images rather than slices, this is not possible due to our current GPU limitations. Accordingly, we designed a two-step training framework for MISPEL which consists of units of 2D encoder and decoder modules for each of the scanners. The 2D network (Figure 2) is trained on axial slices, since this orientation has the highest resolution in our images. More details on MISPEL were provided in (Torbati et al., 2021b) and the code is publicly available^1^.

**Figure 2:**
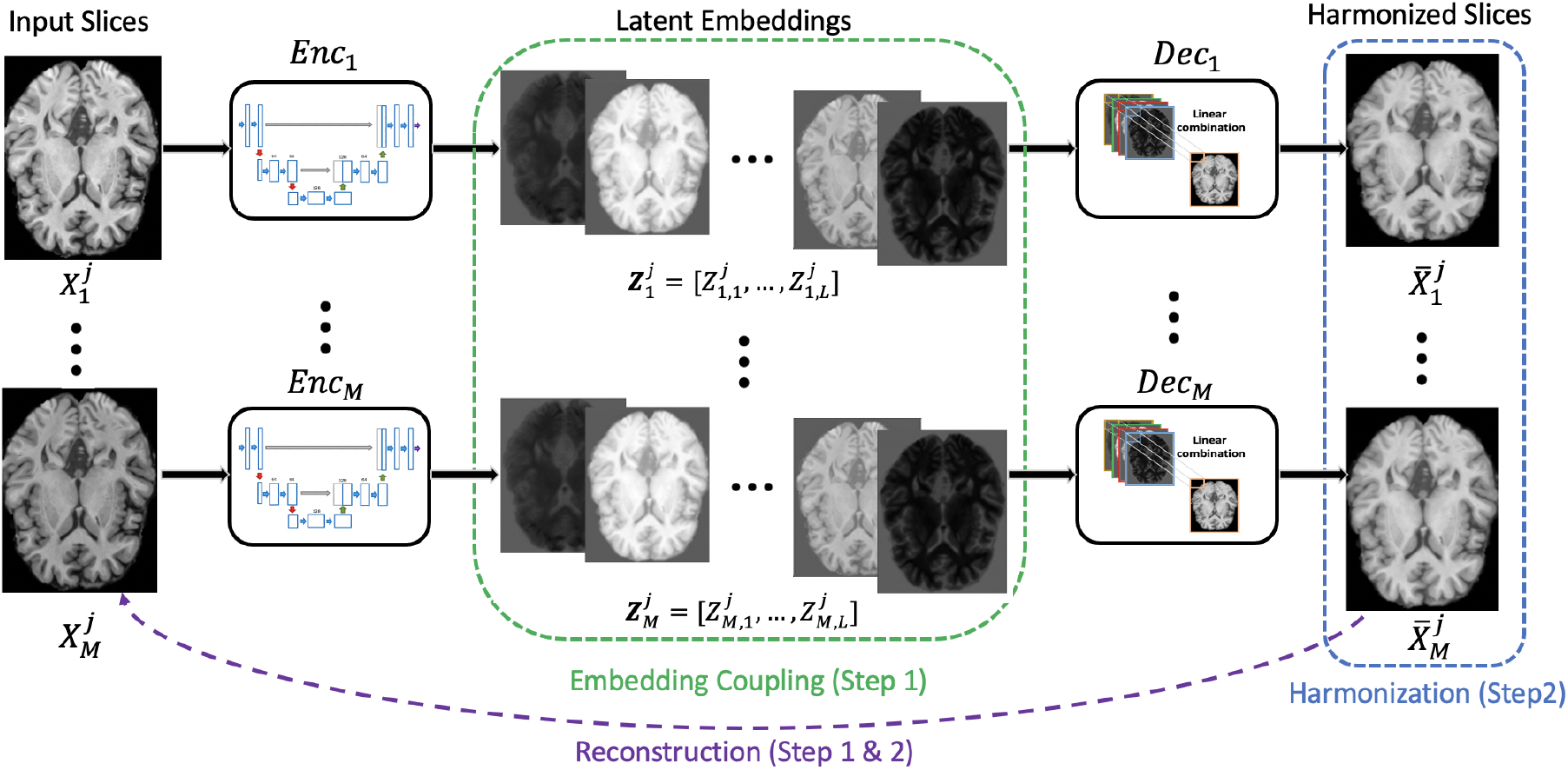
Illustration of MISPEL. For each of the *i* = 1 : *M* scanners and the *j* = 1 : *N* input axial slices, *Enc*_*i*_ (2D U-Net) decomposes the corresponding latent embeddings: 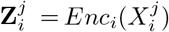). The corresponding *Dec*_*i*_ (linear function) then maps the embeddings to the output: 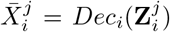. **Step 1** Embedding Learning: *Enc*_*i*_-*Dec*_*i*_ unit reconstructs the input images for each scanner *i*. In this step, *Enc*_*i*=1:*M*_ and *Dec*_*i*=1:*M*_ are updated using the Embedding Coupling loss and the Reconstruction loss. **Step 2** Harmonization: the *Dec*_*i*_ synthesizes the harmonized images for each scanner *i*. In this step, only *Dec*_*i*=1:*M*_ are updated using the Harmonization loss and the Reconstruction loss.

#### 2.3.1 Implementation

We consider *M* scanners for RAW data, i.e., the preprocessed matched images which are registered to the same template space. The axial slices across all RAW scans are combined for a total of *N* slices for each scanner *i, i* = 1 : *M*. The dataset thus consists of 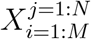, where 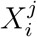 is the slice *j* from scanner *i* and 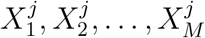 are the matched axial slices. Our goal is to achieve the harmonized axial slices, referred to as 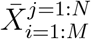, by making them similar across scanners, i.e., achieving 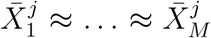, for *j* = 1 : *N*.

For network generalizability and expandability to multiple scanners, MISPEL uses a separate unit of encoders and decoders for each of the scanners. We designed *Enc*_*i*_ (the encoder for scanner *i*) as a 2D U-Net (Ronneberger et al., 2015), which decomposes slice 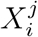 into its set of *L* latent embeddings 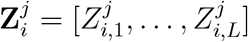. *Dec*_*i*_ is then designed as a linear function combining the components of latent embeddings, 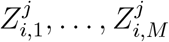, to map 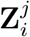 to 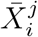.

#### 2.3.2 Network training

Each *Enc*_*i*_-*Dec*_*i*_ unit reconstructs 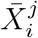 from 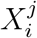 for each scanner *i* and slice *j* and cannot reach harmonization by itself. Thus, we employ another mechanism in order to make the synthesized images, 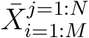, similar across the scanners and achieve harmonization. One way to do that would be to train all *Enc*-*Dec* units to directly impose similarity of matched slices by a loss function. However, this may result in modification of brain structures, as we noticed that even our matched slices which were co-registered in the preprocessing, have small structural differences. Thus, we implemented a two-step training for MISPEL which preserves the brain structure. In Step 1, we first learned the embeddings with structural information, and in Step 2, we harmonized the intensities of embeddings without modifying the structure of the brains.

### Step 1: Embedding Learning

For learning embeddings that could preserve the structural information of the brain, we train the *Enc*-*Dec* units to reconstruct their corresponding input slices. For example, for scanner *i* and slice *j*, the goal for *Enc*_*i*_*Dec*_*i*_ is to achieve 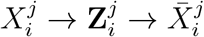, in which 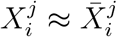. To enforce all units to image reconstruction, we used Reconstruction loss (ℒ _*recon*_). ℒ _*recon*_ should enforce all units to reconstruct their input images. To use this specific Reconstruction loss, we first compute the pixel-wise mean absolute error (MAE) between 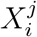 and 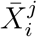 for *i* = 1 : *M* and then sum them over. In addition to this image reconstruction strategy, the *Dec*_*i*_ modules maintain the structural information of the brain by *linearly* combining the embeddings.

Making the latent embeddings similar across scanners will improve the results of harmonization later in Step 2. By this similarity, for example for scanner *i* and slice *j*, the goal is to obtain 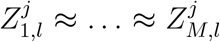, for *l* = 1 : *L*. For enforcing the similarity, we designed the Embedding Coupling loss (*L*_*coup*_) to “couple” the embeddings of the *M* scanners. We first calculated the pixel-wise variance for the *l*th embedding over all *M* scanners. We conducted this step for all *L* embeddings. We then calculated ℒ _*coup*_ as the mean of these variances over all embeddings and their pixels. The loss for Step 1 is then calculated as ℒ _*step*1_ = *λ*_1_ ℒ_*recon*_ + *λ*_2_ ℒ_*coup*_, where *λ*_1_ *>* 0 and *λ*_2_ *>* 0 are the weights. We trained our units of *Enc*-*Dec* for *j* = 1 : *N* slices. The units 13 trained simultaneously for *T*_1_ times.

### Step 2: Harmonization

We continue the training process with Step 2 in which for each scanner *i* and slice *j*, the goal is to achieve 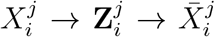. Unlike Step 1, the 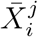 will be the harmonized slice in this step. For harmonizing slices, we froze the encoders during the training and updated just the decoders to synthesize similar matched slices, i.e., achieving 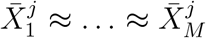. For enforcing the similarity, we used the Harmonization loss (ℒ _*harm*_). We first calculated the MAEs between the images of all unique scanner pairs. ℒ _*harm*_ was then the mean of these MAEs. For example, for slice *j*, the ℒ _*harm*_ is the mean of MAEs for 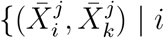, *k* ∈ *{*1, …, *M }* and *i < k }*. In the loss for Step 2, we also incorporate ℒ _*recon*_ to discourage deviation of harmonized images from their originals. Thus, we have ℒ _*step*2_ = *λ*_3_ ℒ _*recon*_ + *λ*_4_ℒ _*harm*_, where *λ*_3_ *>* 0 and *λ*_4_ *>* 0. With ℒ _*step*2_, we trained the decoders of all units for *j* = 1 : *N* slices and repeat it for *T*_2_ times, when ℒ _*step*2_ does not change anymore. By the end of this step, the synthesized images, 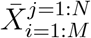, are the desired harmonized ones.

#### 2.3.3 Harmonization practicality

The general harmonization approach for *supervised* methods is to use matched data to learn scanner effects of the scanners for which the matched data were collected (Dewey et al., 2019; Wrobel et al., 2020). Such a trained model could then be used for harmonization of the images taken by any of these scanners. These images do not necessarily need to be matched, and harmonization can be applied to images of each scanner separately. For showing such practicality of MISPEL in harmonization, we conducted a 6-fold cross-validation at the subject level using a 12/3/3 split for training, validation, and testing, respectively. In this manner, the images of validation and test sets are treated as unmatched images and are harmonized individually. Moreover, these images are harmonized by models that have not seen them during their training.

We used RAW images as the input of MISPEL. As explained in Section 2.3.2, we started by training each of the 6 models (i.e. datasets) with Step 1 and then continued with Step 2. For tuning the hyper-parameters of the models, we used the images of the validation sets. In Step 1, we fixed *λ*_1_ = 1 and trained models for *λ*_2_ ∈ *{*0.1, 0.2, 0.3, 0.4, 0.5*}* and *L* ∈ *{*4, 6, 8*}*. We then selected *L* = 6, *λ*_1_ = 1, and *λ*_2_ = 0.3 based on the average of ℒ _*step*1_ values on the validation sets. In Step 2, we fixed the models for *λ*_3_ = 1 and trained the models for *λ*_4_ = *{*1, 2, 3, 4, 5, 6*}*. We ended up having *λ*_3_ = 1 and *λ*_4_ = 4, when we compared results based on the average of the ℒ _*step*2_ values for the validation sets. The training was conducted on NVIDIA RTX5000 for *T*_1_ = 100 and *T*_2_ = 100 with the batch size of 4. For both steps, we used ADAM optimizer (Kingma and Ba, 2014) with a learning rate of 0.01. Training of each model took approximately 200 and 30 minutes for Step 1 and Step 2, respectively.

We then used the tuned models for harmonizing their corresponding test sets. In the next section, we explain that two of our competing methods, WS and RAVEL, were designed to be applied to all images at once. For ease of comparing MISPEL to these methods, we pooled all the MISPEL-harmonized test sets as one harmonized set. This is the dataset that is used in Section 3 for reporting the harmonization performance of MISPEL.

### 2.4 Competing methods

We compared MISPEL with one method of intensity normalization, White Stripe (WS), and two methods of harmonization, RAVEL and CALAMITI. We selected WS and RAVEL as they (1) are widely applied to MRI neuroimaging data, (2) can be applied to multiple (more than two) scanners, and (3) do not require specifications of a *target* scanner. We considered CALAMITI as our main competing method since it can be slightly modified and applied to matched data, and could be regarded as one of the state-of-the-art methods in harmonization. We emphasize that determining the ultimate state-of-the-art harmonization method is not trivial as harmonization lacks standardized evaluation criteria.

**White Stripe (WS)** is an individual-level intensity normalization method for removing discrepancy of intensities across subjects within tissue types (Shinohara et al., 2014b). It first extracts the normal-appearing white matter voxels of the image and estimates moments of their intensity distribution. It then uses these moments in the z-score transformation for normalizing the voxels of all brain tissue types.

**RAVEL** is an intensity normalization and harmonization framework (Fortin et al., 2016). It initializes with a WS normalization step and then applies a voxel-wise harmonization strategy to images. In the harmonization strategy, RAVEL first estimates the components of scanner effects by applying singular value decomposition to cerebrospinal fluid (CSF) voxels of images. These voxels are known to be unassociated with disease status and clinical covariates and are representative of scanner effects. RAVEL then uses these voxels to estimate scanner effects and harmonizes the images by removing the estimated scanner effects from the voxel intensities. Throughout the estimation of the scanner effects, we considered the status of the subjects (cognitively normal with low or high degree of SVD) as the biological/clinical covariates. We also set the components of scanner effects to 1, as suggested in the original work (Fortin et al., 2016). For further details on the biological/clinical covariates and components of scanner effects, see Algorithm 1 in (Torbati et al., 2021a).

**CALAMITI** is an unsupervised deep-learning method for harmonizing multiscanner inter-modality paired dataset (Zuo et al., 2021). It is a domain adaptation approach mapping the images of scanners to the domain of a *target* scanner. Intermodality paired dataset consists of images of two predetermined modalities taken from one individual on the *same* scanner with a short time gap. This dataset can have paired images of multiple scanners. For simplicity, we refer to these images as *paired* in the description of this method. CALAMITI should be first trained on paired images of two scanners, one of which should be the *target* scanner. It could then be fine-tuned to map images of other scanners to the target domain. During the training, CALAMITI first gets the paired images as inputs and generates a disentangled representation that captures the mutual scanner-invariant anatomical information (*β*) of images as well as the contrast information (*θ*)s of their modalities and scanner, and then synthesizes the input paired images using their generated mutual *β* and *θ*s. For harmonizing an input image, the trained model is used to generate the *β* of the image and *θ* of one random image from the target scanner. The model then synthesizes the harmonized image using these two components.

We used CALAMITI as a supervised method by simply training it on our *interscanner* paired data. Like MISPEL, we used the 6-fold cross-validation strategy for training and testing the models. We also pooled the harmonized test sets to have one set of data to report the harmonization performance of CALAMITI in Section 3. Following its original paper, we went through one step of normalization and trained CALAMITI using the WS-normalized RAW images. Instead of conducting finetuning, we went for a simpler approach and trained 3 individual models to map GE, Philips, and SiemensP to SiemensT. We used the same machines used for MISPEL and trained CALAMITI with the hyper-parameters reported in its original paper. For being comparable and fair to other methods, we trained CALAMITI on 2D axial slices and skipped its super-resolution preprocessing step and post-harmonization slice-to-slice consistency enhancement step.

Among the competing methods, we regard CALAMITI as a state-of-the-art harmonization method to compare against MISPEL, and we emphasize that WS and RAVEL were not designed to use matched data in their technical variability removal process. Specifically, WS is an intensity normalization method, which does not account for scanner information. However, it is beneficial to study scanner effects and harmonization on the WS-normalized data to emphasize the importance of harmonization for neuroimaging data. On the other hand, RAVEL was designed to remove the inter-subject technical variability of images after intensity normalization. Although RAVEL does not account for scanner information either, scanner effects may appear in the singular value decomposition component extracted individually for each of the subjects from their CSF tissue in this framework. As such, we regard RAVEL as a normalization and harmonization framework that can be compared to CALAMITI and MISPEL to evaluate the advantages of using and accounting for matched data in harmonization methodology.

### 2.5 Data analysis

A harmonization method is expected to remove scanner effects while preserving the biological variables of interest in the data. In our specific matched dataset, the matched images are assumed to be biologically identical but differ entirely due to scanner differences. Thus, the scanner effects can be estimated as dissimilarity of the matched images, and removing the scanner effects can be regarded as increasing their similarity. We investigated the similarity and dissimilarity of matched images using four evaluation criteria: (1) image similarity, (2) GM-WM contrast similarity, (3) volumetric and segmentation similarity, and (4) biological similarity. We also selected SVD as the clinical signal of interest in our data and investigated whether we could preserve or even enhance the SVD group differences in our data after harmonization. We performed our evaluation metrics for all five methods: RAW, White Stripe, RAVEL, CALAMITI, and MISPEL. The entire matched dataset was used in evaluating each method unless otherwise mentioned. Many of our evaluation metrics require pairwise image-to-image comparison, for which we considered all possible combinations of *scanner pairs*: *{*(GE, Philips), (GE, SiemensP), (GE, SiemensT), (Philips, SiemensP), (Philips, SiemensT), and (SiemensP, SiemensT)*}*. Throughout this manuscript, the two matched images of each scanner pair are referred to as *paired* images. To determine the statistical significance of any comparisons, we used paired *t* -test with *p <* 0.05 denoting the significance.

Scanner effects could appear as contrast dissimilarity across images of different scanners (Dewey et al., 2019, 2020; Liu et al., 2021). More specifically, such dissimilarity could appear as tissue-specific contrast differences in images (Meyer et al., 2019). We, therefore, assessed scanner effects and evaluated harmonization using an **image similarity** metric to measure the similarity of cross-scanner images in their appearance, as well as a **GM-WM contrast similarity** metric to assess the tissue contrast similarity of images.

We first investigated the **image similarity**. For this, we assessed the visual quality of the matched *slices* for all methods. We also quantified the similarity of *all* paired images using the structural similarity index measure (SSIM). SSIM is a pairwise metric that compares two images in terms of their luminance, contrast, and structure. A harmonization method is expected to increase the visual and structural similarity of paired images.

Second, we investigated the **GM-WM contrast similarity** of images. The GM-WM contrast can highly influence the quality of segmentation methods, and increased contrast is expected to result in more accurate segmentation. The GMWM contrast of an image can be estimated as the separability of its histograms of GM and WM voxels. This separability was used as the classification of GM and WM voxels of an image in (Torbati et al., 2021a) and reported as the area under the receiver operating characteristic (AUROC) values, with AUROC = 1 denoting perfect classification (complete separation of histograms) and AUROC = 0.5 showing random classification (complete overlap of histograms). For calculating these AUROC values, we conducted the procedure explained in (Torbati et al., 2021a) for each of the images. We first labeled GM and WM voxels of the image using the tissue mask in the EveTemplate package (Oishi et al., 2009). We then classified these voxels using intensity thresholds selected from the range of intensity values of the GM and WM voxels. Lastly, we formed the AUC curve of the image using the result of each classification. A harmonization method is expected to increase the GM-WM contrast similarity.

Third, we investigated the **volumetric and segmentation similarity** criterion for images. The most practical benefit of harmonization is to enable unbiased multiscanner neuroimaging analyses with minimal scanner effects. Tissue-specific regional neuroimaging measures are the basis of these analyses, and therefore, the volumetric and segmentation similarity of these measures across paired images is a crucial metric for evaluating harmonization. We segmented and measured the volumes of the two brain tissue types: GM and WM. We then analyzed the similarity of *each of* these two tissue types *separately* in four ways: (1) volume distributions, (2) volumetric bias, (3) volumetric variance, and (4) segmentation overlap. For volumetric distributions, we compared the distributions of volumes of each tissue type across their four scanners. These plots show the harmonization performance of methods as the similarity of the distributions of their harmonized measures across scanners. Most of the metrics used in the three other criteria are pairwise comparisons, thus we applied them *separately* to all of the 6 *scanner pairs*. Volumetric bias and variance are two metrics assessing the similarity of measures across scanners in two different ways. For volumetric bias, we calculated the absolute differences between volumes of paired images of each scanner pair and evaluated the harmonization based on the mean of these differences over all individuals of the scanner pair. We used root-mean-square deviation (RMSD) for estimating the volumetric variance of paired images of all individuals within each scanner pair. RMSD of a scanner pair denotes the deviation of volumes of one scanner from that of the other scanner. Lastly, we used Dice similarity score (DSC) to estimate the overlap of tissue segmentation of paired images of each scanner pair. The mean of these DSC values over paired images of all subjects was used as an evaluation metric for harmonization. A harmonization method is expected to result in (1) similar distribution of volumes across scanners, (2) minimal (ideally zero) bias, (3) minimal (ideally zero) variance, and (4) maximal (ideally complete) segmentation overlap; for both tissue types and all scanner pairs.

We conducted the volumetric and segmentation similarity evaluation using two segmentation tools: (1) FSL FAST (version 6.0.3) (Zhang et al., 2001), and (2) segmentation in Statistical Parametric Mapping (SPM12) (Ashburner and Friston, 2005). These frameworks are widely used for tissue segmentation in neuroimaging studies, however, the results of these two segmentation algorithms could have moderate to large differences (Tudorascu et al., 2016). We, therefore, assessed volumes from each segmentation tool independently. Originally, the output of WS, RAVEL, CALAMITI, and MISPEL methods were images in template space, as all methods used RAW images as input. The RAW images were non-linearly registered to a T1-w image atlas (Oishi et al., 2009) in the preprocessing step, Section 2.2. Using their inverse transformations, processed images of all methods were transferred to their native space and then used as inputs of the two segmentation tools for tissue volume extraction and then volumetric similarity evaluation. On the other hand, for having a meaningful tissue segmentation overlap, segmentations and accordingly their images should remain in their template space. Thus, we also ran FSL and SPM frameworks on the template-space images to generate the segmentations and then evaluate the segmentation overlap similarity. For all runs of the segmentation frameworks, we set the tissue class probability thresholds to 0.8.

Fourth, we investigated the **biological similarity** of images using biomarkers of Alzheimer’s disease (AD). We studied the bias (mean of cross-scanner absolute differences) and variance (RMSD) for these biomarkers. For bias, we calculated the cross-scanner absolute differences of all scanner pairs and reported their mean (SD). For variance, we calculated the mean of RMSDs across all scanner pairs. We report these metrics for all 5 methods and all biomarkers of AD. As biomarkers of AD, we investigated cortical thickness measures of the entorhinal and inferior temporal cortices, as well as volume measures of the hippocampus and amygdala. These summary measures are the sum of measures over both hemispheres, and they were extracted using FreeSurfer 7.1.1 (FS) (Fischl, 2012). These regions have previously been found to be most relevant to AD (Schwarz et al., 2016). We extracted these measures across all harmonization methods for 17 of the 18 total subjects. RAVEL-harmonized images of a single subject failed FS segmentation due to an error in the corpus callosum segmentation step. Thus, for a fair comparison across methods, we omitted this subject from the experiments on biomarkers of AD. We also skipped skull stripping and bias correction steps in the FS processing pipeline, as RAW images had already gone through skull-stripping and N4 bias correction during image preprocessing (Section 2.2).

Fifth and last, we investigated whether each harmonization method **preserved or even enhanced a biological/clinical signal of interest** in our matched data. We selected SVD as our clinical signal of interest and investigated the effect size between two groups of low and high SVD in our data. For this experiment, we calculated Cohen’s d effect sizes of the two SVD groups for each of our FS-derived biomarkers of AD individually. For each of the biomarkers, we calculated the size effects of the scanners separately and reported the mean (SD) of these values across scanners. A harmonization method is expected to not deteriorate the effect sizes of groups after harmonization.

## 3 Results

In this section, we report our evaluation criteria on RAW, WS-normalized, RAVEL-, CALAMITI-, and MISPEL-harmonized images. For a more convenient comparison with RAW, WS and RAVEL, we pooled harmonized images of each of CALAMITI and MISPEL as one dataset.

### 3.1 Image similarity

The similarity of images across normalization and harmonization methods is depicted in Figures 3 and 4. Visual assessment of processed images in Figure 3 revealed that (1) scanner effects are present in the matched RAW images and appear most significantly as differences in image contrast, (2) White Stripe made matched images more similar, but at the expense of decreased contrast, (3) RAVEL improved upon WS by increasing contrast relative to WS-normalized images, (4) CALAMITI improved similarity of the matched images by adapting contrast across all scanners to that of the RAW SiemensT, and (5) MISPEL improved the similarity of images similarly to CALAMITI but visually smoothed images to some extent.

**Figure 3:**
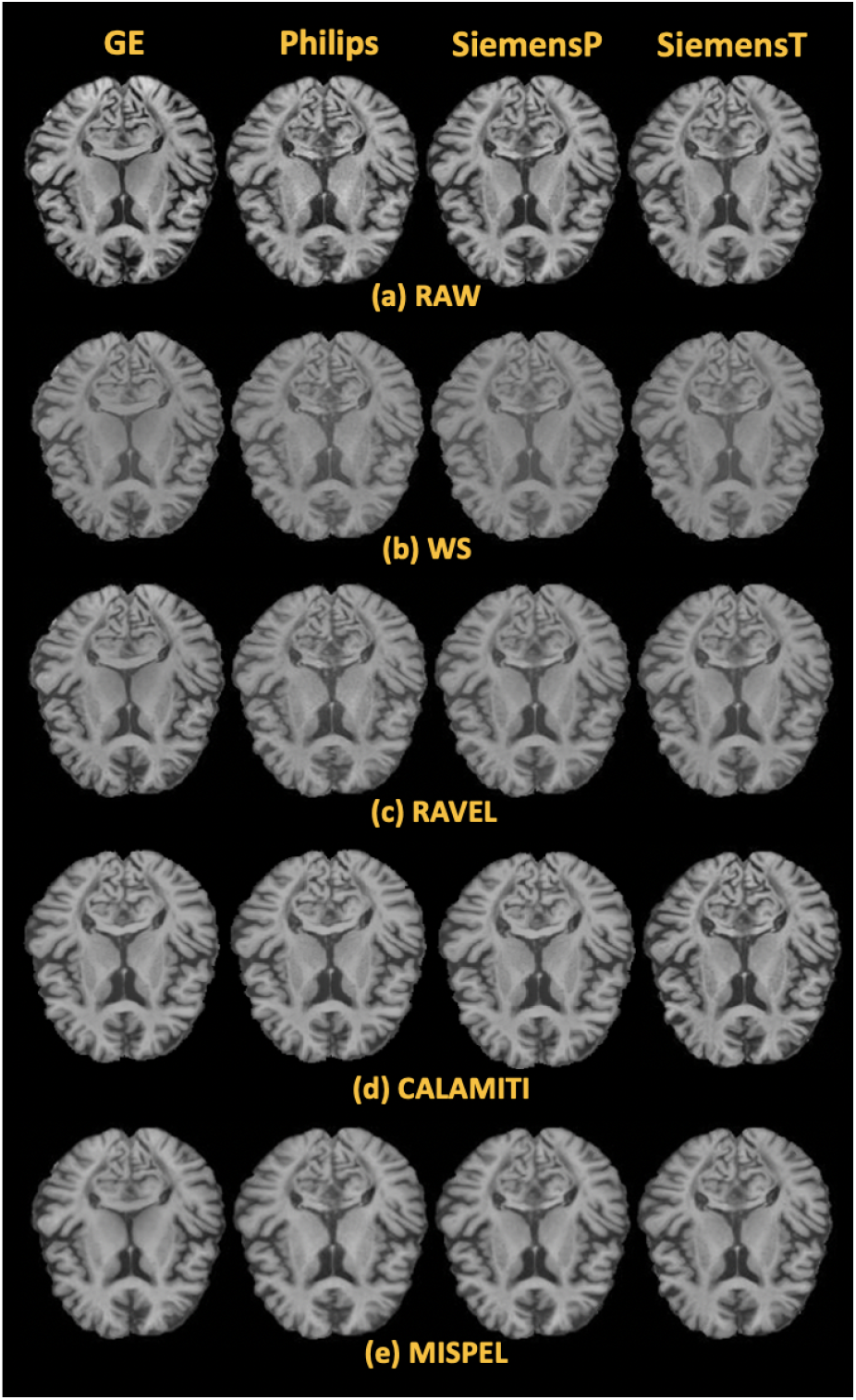
Visual assessment of matched images of a slice. Rows and columns correspond to methods and scanners respectively. All four methods: WS, RAVEL, CALAMITI, and MISPEL made the matched slices of RAW more similar, with CALAMITI and MISPEL preserving their contrast the most.

**Figure 4:**
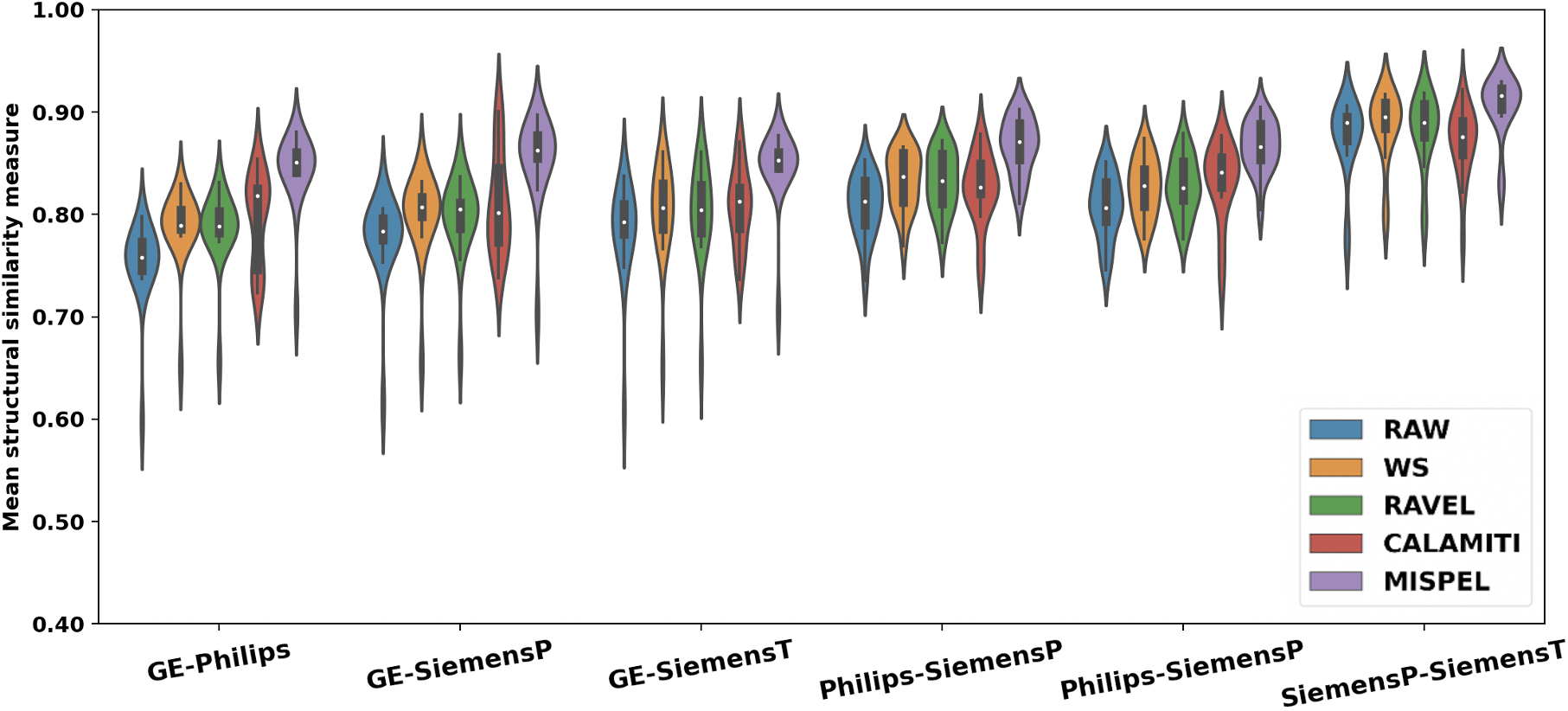
Structural similarity index measures (SSIM) for the matched dataset. The SSIM distributions of images of scanner pairs were depicted as violin plots for each of the methods. A harmonization method is expected to have the highest mean of SSIM. MISPEL improved SSIMs of RAW and outperformed the other three methods. For interpretation of the references to color in this figure legend, the reader is referred to the web version of this article.

For a quantitative understanding of similarity of images, we explored the SSIM distribution of the matched images of all subjects for the 6 *scanner pairs* enumerated in Section 2.5. These distributions are depicted as violin plots for the five methods: RAW, WS, RAVEL, CALAMITI, and MISPEL in Figure 4. The violin plots with the smallest SSIM mean belong to RAW, indicating scanner effects exist in our matched dataset as dissimilarity of images. Scanner pairs including GE have long-tailed distributions, which indicates that GE images are most dissimilar to others. Moreover, the SiemensP-SiemensT scanner pair had the largest SSIM mean, indicating that these two are the most similar scanners.

We observed that WS, RAVEL, CALAMITI, and MISPEL improved SSIM of RAW for all of its scanner pairs, except for CALAMITI for the SiemensP-SiemensT scanner pair. Lastly, we observed that MISPEL outperformed the other three methods. All comparisons were statistically significant (assessed using paired *t* -tests), except for CALAMITI for the Philips-SiemensP and SiemensP-SiemensT pairs.

### 3.2 GM-WM contrast similarity

We quantified the GM-WM contrast of an image using the AUROC values denoting the separation of histograms GM and WM voxel intensities. High AUROC indicates higher contrast, with 100% the highest. In Figure 5, we depicted the spaghetti plots of AUROC values of images of all subjects across the four scanners. A harmonization method is expected to (1) make the AUROC of matched images similar, i.e., results in overlapped lines, and (2) not deteriorate the AUROC of images.

**Figure 5:**
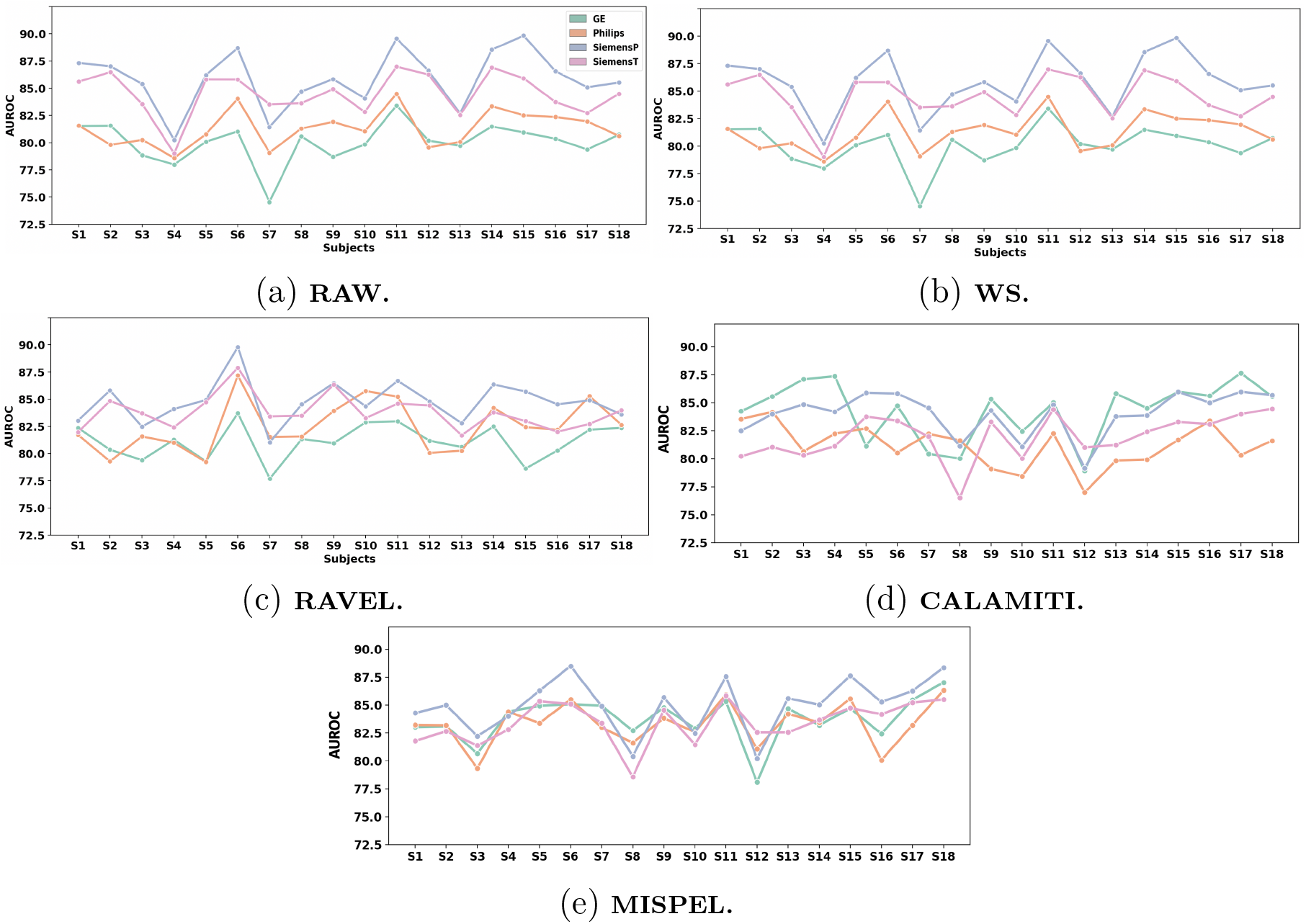
GM-WM contrast spaghetti plots. The GM-WM contrast was estimated as AUROC values and was depicted for images of all subjects as spaghetti plots. In these plots, each line corresponds to one scanner. A harmonization method that performs well should show overlap of the lines. Plots showed that MISPEL outperformed WS, RAVEL, and CALAMITI with the highest overlapped of the lines. For interpretation of the references to color in this figure legend, the reader is referred to the web version of this article.

Figure 5a shows that scanner effects exist in RAW data and appeared as dissimilarity of GM-WM contrast in matched dataset, i.e., distant lines in this plot. Figure 5b shows that WS does not change AUROCs of RAW. On the other hand, Figures 5c, 5d, and 5e show respectively that RAVEL, CALAMITI, and MISPEL resulted in more overlapped lines, with MISPEL having the highest overlap.

Figure 6 shows the bar plots indicating the mean AUROC of images of each scanner. MISPEL is the only method that increased the mean AUROC of RAW images for all scanners. We also observed that: (1) WS did not change the mean AUROC value of RAW, (2) RAVEL improved the contrast for GE and Philips, but made it worse for SiemensP and SiemensT, and (3) CALAMITI improved the mean AUROC of GE and Philips and did not affect that of other scanners. In addition to these results, MISPEL seems to be the most successful method in bringing the mean AUROC of the scanners closer to each other. In summary, we show that MISPEL is the only method that satisfied both harmonization criteria determined for GM-WM contrast similarity.

**Figure 6:**
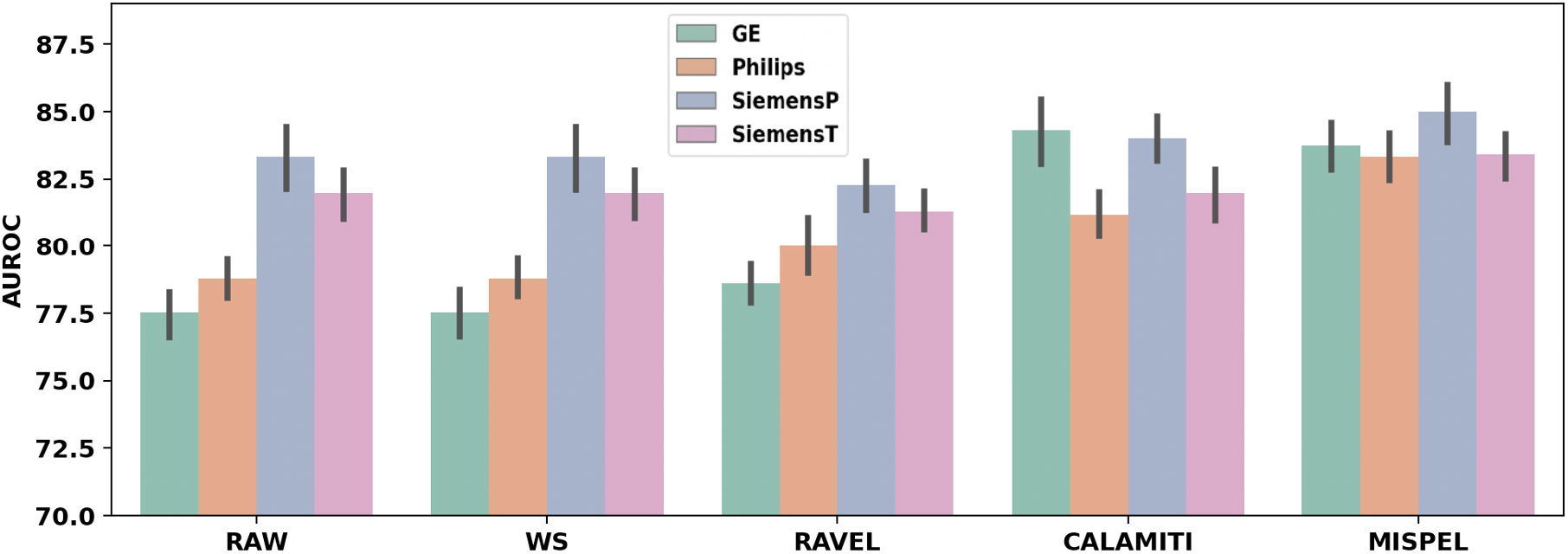
GM-WM contrast bar plots. Each bar indicates the mean AUROC of images of each scanner, with error bars denoting the standard deviation for each method. A harmonization method is expected to not deteriorate the GM-WM contrast of images. Plots show that MISPEL outperformed WS, RAVEL, and CALAMITI reflected in the similarity of the boxplots. For interpretation of the references to color in this figure legend, the reader is referred to the web version of this article.

### 3.3 Volumetric and segmentation similarity

We estimated the volumetric and segmentation similarity of GM and WM tissue types based on four criteria: (1) volume distributions, (2) volumetric bias, (3) volumetric variance, and (4) segmentation overlap. We performed our evaluation for FSL and SPM segmentation frameworks and expected the harmonization methods to result in: (1) similar volume distributions across scanners, (2) minimal bias, (3) minimal variance, and (4) maximal segmentation overlap; for both tissue types and both segmentation frameworks.

#### 3.3.1 Volume distributions

Figure 7 shows boxplots of volumes of the two tissue types, GM and WM, across the four scanners for all five methods, with Figures 7a and 7b depicting these boxplots for volumes extracted by FSL and SPM frameworks, respectively. Plots in Figure 7a showed that scanner effects exist in the matched volumes derived through FSL and appeared as dissimilar boxplots for RAW across scanners. When compared to RAW, WS and RAVEL resulted in more dissimilar boxplots for FSL-derived volumes of both GM and WM. On the other hand, we noticed that the use of CALAMITI and MISPEL helped towards harmonization of data. CALAMITI made GE and Philips more similar to SiemensP and SiemensT for both GM and WM, but increased variance for distributions of all scanners for WM. Similarly, MISPEL made GE more similar to SiemensP and SiemensT for both GM and WM volumes. Figure 7b showed that scanner effects exist in RAW volumes extracted by SPM too. Our normalization and harmonization methods though resulted in relatively minor changes in SPM-derived GM and WM volumes, with CALAMITI and MISPEL showing the most noticeable changes. Both CALAMITI and MISPEL made Philips closer to SiemensP and SiemensT for GM volumes. They also made GE closer to these two scanners for WM.

**Figure 7:**
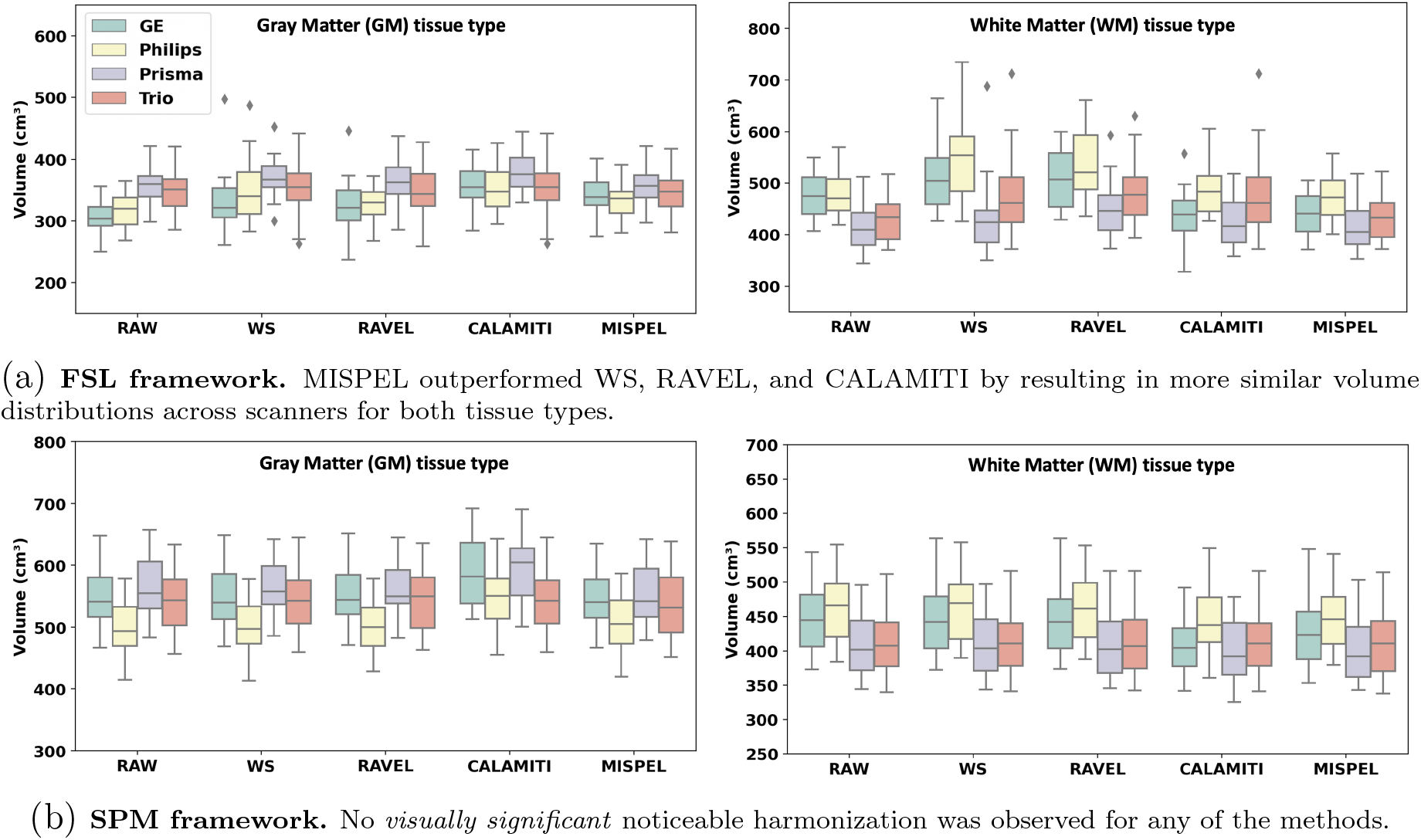
Volume distribution boxplots. Boxplots denote the volume distribution of GM and WM tissue types for images of each scanner. These boxplots were depicted for all five methods and explored for two segmentation frameworks: (a) FSL and (b) SPM. A harmonization method is expected to result in similar distributions of volumes across scanners. For interpretation of the references to color in this figure legend, the reader is referred to the web version of this article.

In summary, MISPEL and CALAMITI outperformed WS and RAVEL in harmonizing FSL-derived volumes and none of the methods resulted in *visually significant* assessed harmonization for the SPM-derived volumes, when volumetric distribution similarity of *both* GM and WM volumes were used as the evaluation metric. Results for the statistical assessment of harmonization of FSL- and SPM-derived GM and WM volumes are presented in the next section.

#### 3.3.2 Volumetric bias

Table 2 shows mean and standard deviation (SD) of cross-scanner absolute differences of all paired volumes in each scanner pair. We calculated these statistics for volumes of GM and WM tissue types extracted using FSL and SPM segmentation frameworks, for all five methods. We also presented the distributions of these differences as violin plots in Figure 8. Using paired *t* -test, we compared each of these distributions to their equivalent distributions in RAW.

**Table 2:** Mean absolute differences for GM and WM volumes. Mean (SD) of cross-scanner absolute differences of volumes for all scanner pairs and all methods. The volumes are for GM and WM tissue types and were extracted through two segmentation frameworks: FSL and SPM. A harmonization method that works is expected to have low values of mean and SD for all paired scanners. MISPEL outperformed WS, RAVEL, and CALAMITI by having the largest number of smallest means and not significantly increasing the values of SD, for both FSL and SPM. The distributions with the smallest means are in bold. Also, the distributions that showed statistically significant t-statistics when compared to RAW were marked by *.

| Framework | Tissue | Method | Mean (SD) of Volumetric Absolute Differences for Paired Dataset |  |  |  |  |  |
| --- | --- | --- | --- | --- | --- | --- | --- | --- |
|  |  |  | GE-Philips | GE-SiemensP | GE-SiemensT | Philips-SiemensP | Philips-SiemensT | SiemensP-SiemensT |
| FSL | GM | RAW | 19.82 (9.10) | 55.84 (16.54) | 46.53 (16.94) | 39.70 (15.28) | 29.00 (15.75) | 12.14 (9.17) |
|  |  | WS | 43.53 (56.27) | 56.66 (29.56) | 46.34 (31.84) | 49.31 (40.21) | 43.09 (49.92) | 18.00 (15.37) |
|  |  | RAVEL | 27.53 (32.28) | 52.88 (16.60) | 39.22 (22.41) | 38.87 (17.79) | 24.65 (20.50) | 17.53 (9.35) |
|  |  | CALAMITI | 32.29 (21.87)* | 26.07 (33.88)* | 32.18 (31.05) | 28.06 (16.00)* | 26.07 (17.98) | 26.02 (26.47)* |
|  |  | MISPEL | **11.06** (17.76) | **20.57** (10.96)* | **14.83** (14.21)* | **20.74** (10.20)* | **10.97** (9.98)* | **11.98** (7.96) |
|  | WM | RAW | **15.39** (11.29) | 59.59 (20.92) | 42.45 (18.37) | 67.30 (13.41) | 50.16 (16.43) | 17.89 (15.48) |
|  |  | WS | 46.99 (54.09)* | 100.63 (64.18)* | 71.35 (47.72)* | 119.73 (79.23)* | 81.41 (41.29)* | 41.03 (50.11) |
|  |  | RAVEL | 43.95 (36.46)* | 65.02 (37.96) | 42.42 (34.98) | 89.59 (49.34) | 57.60 (23.77) | 32.18 (39.53) |
|  |  | CALAMITI | 61.39 (37.88)* | 28.57 (23.69)* | 59.84 (64.36) | 65.60 (24.08) | 68.64 (46.78) | 58.07 (74.88)* |
|  |  | MISPEL | 38.68 (14.17)* | **25.19** (18.80)* | **15.76** (15.63)* | **58.26** (13.05)* | **43.42** (11.93)* | **15.35** (14.00) |
| SPM | GM | RAW | 48.22 (20.82) | 23.45 (12.67) | 19.37 (11.23) | 63.57 (15.90) | 44.65 (16.94) | 19.86 (13.77) |
|  |  | WS | 48.60 (21.35) | 21.75 (13.15) | **14.94** (12.70) | 65.72 (15.54) | 46.84 (18.99) | 19.46 (13.53) |
|  |  | RAVEL | 46.12 (22.48) | **10.44** (7.57)* | 15.22 (9.77) | 53.82 (18.48)* | 39.14 (20.99) | 15.26 (12.85)* |
|  |  | CALAMITI | 42.22 (36.06) | 37.85 (28.09) | 49.62 (41.39)* | 46.81 (23.06)* | **28.44** (19.67)* | 51.51 (34.64)* |
|  |  | MISPEL | **41.80** (15.26) | 16.05 (9.05)* | 17.40 (13.74) | **42.23** (15.10)* | 30.13 (17.91)* | **14.67** (10.15) |
|  | WM | RAW | 21.06 (15.98) | 40.40 (18.08) | 35.45 (20.87) | 53.16 (11.74) | 48.22 (12.32) | 9.06 (7.31) |
|  |  | WS | 25.97 (20.29)* | 40.18 (23.46) | 34.18 (27.71) | 54.43 (11.43) | 48.80 (13.16) | 9.69 (7.53) |
|  |  | RAVEL | 22.49 (15.69) | 35.60 (19.36)* | 34.02 (21.03) | 47.64 (10.95)* | 46.48 (12.14) | **8.41** (8.00) |
|  |  | CALAMITI | 40.14 (21.52)* | **16.24** (9.80)* | **20.13** (18.07)* | 49.31 (16.99) | **34.67** (16.21)* | 20.21 (17.92)* |
|  |  | MISPEL | **19.45** (14.90) | 27.70 (14.53)* | 20.46 (16.85)* | **43.75** (11.00)* | 35.58 (12.31)* | 14.05 (7.32)* |

**Figure 8:**
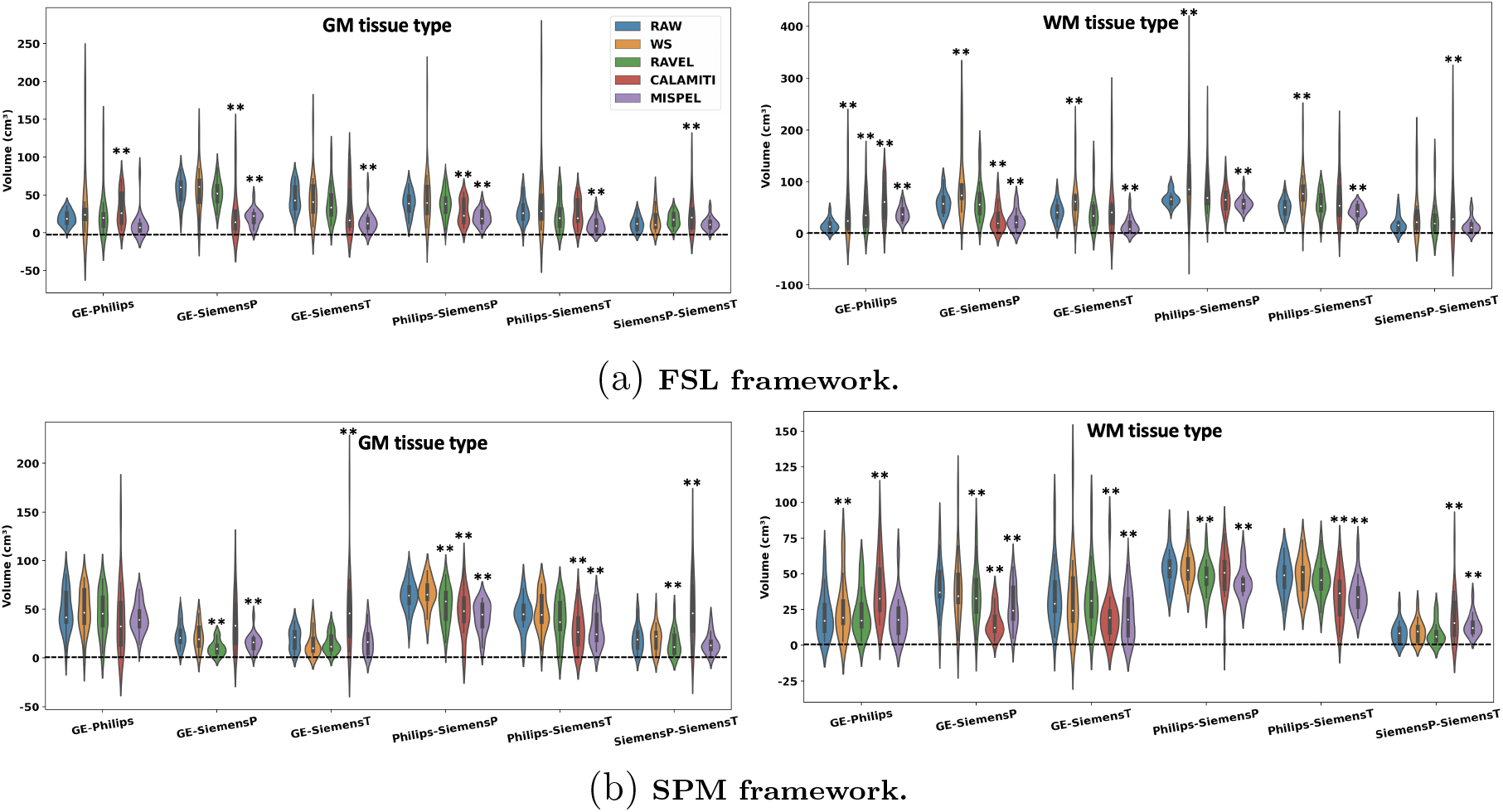
Absolute difference violin plots. The distributions of absolute differences of paired volumes as violin plots for all scanner pairs. The volumes are for GM and WM tissue types and extracted using two segmentation frameworks: (a) FSL and (b) SPM. A harmonization method is expected to result in short and fat (wide) violins, with mean values centered at zero. MISPEL outperformed WS, RAVEL, and CALAMITI by having largest number of these violin plots for both FSL and SPM. The distributions that showed statistically significant t-statistics when compared to RAW were marked by **. For interpretation of the references to color in this figure legend, the reader is referred to the web version of this article.

A harmonization method is expected to result in minimal (ideally zero) mean of absolute differences (bias), with no major increase in SD of the differences. The SD values indicate the consistency of harmonization across subjects. A harmonization method should harmonize images of all subjects to a comparable degree, and thus should not increase the SDs drastically. Likewise visually, the violin plots in Figure 8 for harmonized images are expected to be centered as close as possible to zero.

We observed that scanner effects exist in the RAW volumes extracted through FSL framework and appeared for all scanner pairs as non-zero bias values. We also observed that MISPEL resulted in the largest number of smallest biases for FSL-derived volumes, when compared to the other three methods. This number was 11 out of a total of 12 cases, which are the 6 scanner pairs of the 2 tissue types. 8 out of these 11 biases were significantly different than their equivalents in RAW. Moreover, we noticed that MISPEL did not significantly increase the SD of distributions, just 2 increases out of 12, in which only the SD of GM for the GE-Philips pair had a major increase. On the other hand, WS, RAVEL, and CALAMITI showed increases in SD of differences for all 12 distributions, with WS showing the most drastic increases (Figure 8). In general, RAVEL and CALAMITI harmonized FSL-derived volumes to some extent. Compared to RAW, RAVEL resulted in 5 decreased biases and CALAMITI resulted in 6 decreases. However, CALAMITI also resulted in drastically increased biases for the WM volumes of 5 of the scanner pairs (Figure 8a).

Results of RAW volumes extracted by SPM show that SPM is also sensitive to scanner effects. MISPELand CALAMITI decreased bias for 11 and 7 cases, respectively. They resulted in the largest numbers of smallest biases for SPM: 5 and 4 out of 12 cases for MISPEL and CALAMITI, respectively. Among these cases, 2 for MISPEL and all for CALAMITI showed statistical differences when compared to RAW. On the other hand, CALAMITI increased SD for 8 out of 12 cases, while other methods did not show any major increases. This can be observed in Table 2 as well as Figure 8b. WS and RAVEL harmonized the SPM-derived volumes to some extent by decreasing the biases of 5 and 11 cases, respectively. They also resulted in a few smallest biases: 1 case for WS and 2 cases for RAVEL.

Summarizing Table 2 and Figure 8, we observed that MISPEL outperformed WS, RAVEL, and CALAMITI when FSL and SPM were used for extracting volumes and volumetric bias and SD of differences were used as harmonization evaluation metrics.

#### 3.3.3 Volumetric variance

Figure 9 shows bar plots that indicate the RMSD of paired volumes in each of the scanner pairs. We calculated these values for volumes of GM and WM tissue types and depicted them for all five methods. Figure 9 contains these sets of bar plots for volumes extracted through FSL and SPM frameworks in Figures 9a and 9b, respectively. Ideal harmonization would result in a zero RMSD for each scanner pair.

**Figure 9:**
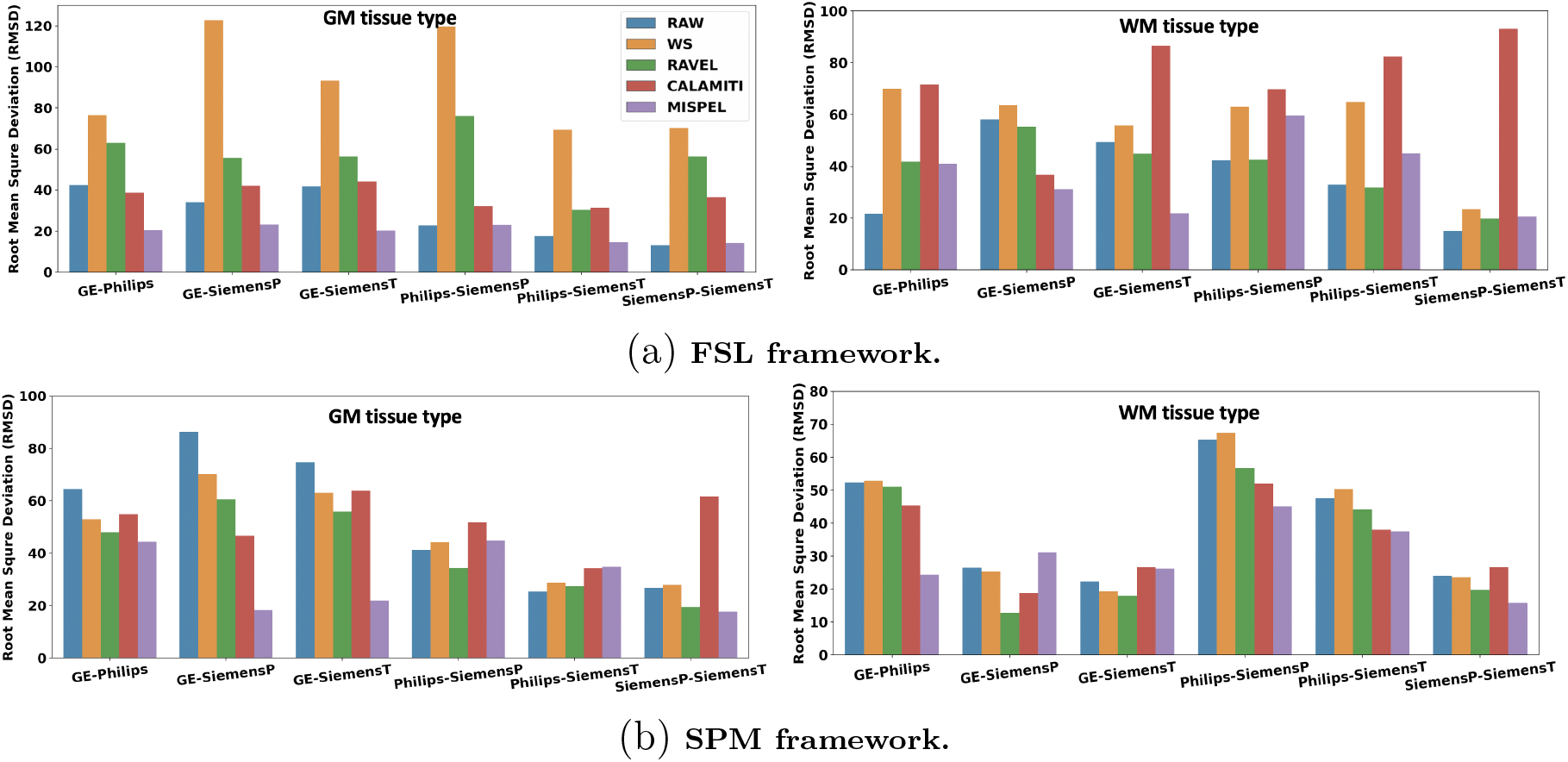
Root-mean-square deviation (RMSD) bar plots for GM and WM volumes. Bar plots indicate the RMSD of paired volumes in scanner pairs. These values were calculated for volumes of GM and WM tissue types and depicted for all five methods. These set of bar plots were depicted for volumes extracted through two segmentation frameworks: (a) FSL and (b) SPM. A harmonization method is expected to lower values of RMSDs. MISPEL outperformed WS, RAVEL, and CALAMITI by having the largest number of smallest RMSD values for volumes of both FSL and SPM. For interpretation of the references to color in this figure legend, the reader is referred to the web version of this article.

We observed that scanner effects exist in RAW volumes for both segmentation frameworks and appeared as non-zero RMSD values. Also, MISPEL outperformed WS, RAVEL, and CALAMITI, showing the smallest RMSD values: 6 and 8 out of 12 cases for FSL and SPM, respectively. These statistics are 0 and 1 for CALAMITI as well as 0 and 3 for RAVEL. We also observed that WS did not improve the RMSD values of any 12 scanner pairs for FSL, when compared to RAW. However, it performed better for SPM by decreasing the number of worse cases to 6. MISPEL, CALAMITI, and RAVEL deteriorated some of the RMSDs too. Among these methods, MISPEL deteriorated the least number of cases, 4 for each of the FSL- and SPM-derived volumes.

In summary, we observed that MISPEL outperformed WS, RAVEL, and CALAMITI when FSL and SPM were used for deriving volumes and volumetric variance was used as the harmonization evaluation metric.

#### 3.3.4 Segmentation overlap

Figure 10 shows bar plots that indicate the mean DSC of all paired segmentations in each scanner pair. We calculated the means of DSCs for segmentations of GM and WM tissue types and depicted them for all five methods. Figure 10 contains these sets of bar plots for segmentations extracted through FSL and SPM frameworks in Figures 10a and 10b, respectively. DSC shows the overlap of two paired segmentations. A good harmonization method would result in an increased mean of DCSs for all scanner pairs, with 1 indicating the highest.

**Figure 10:**
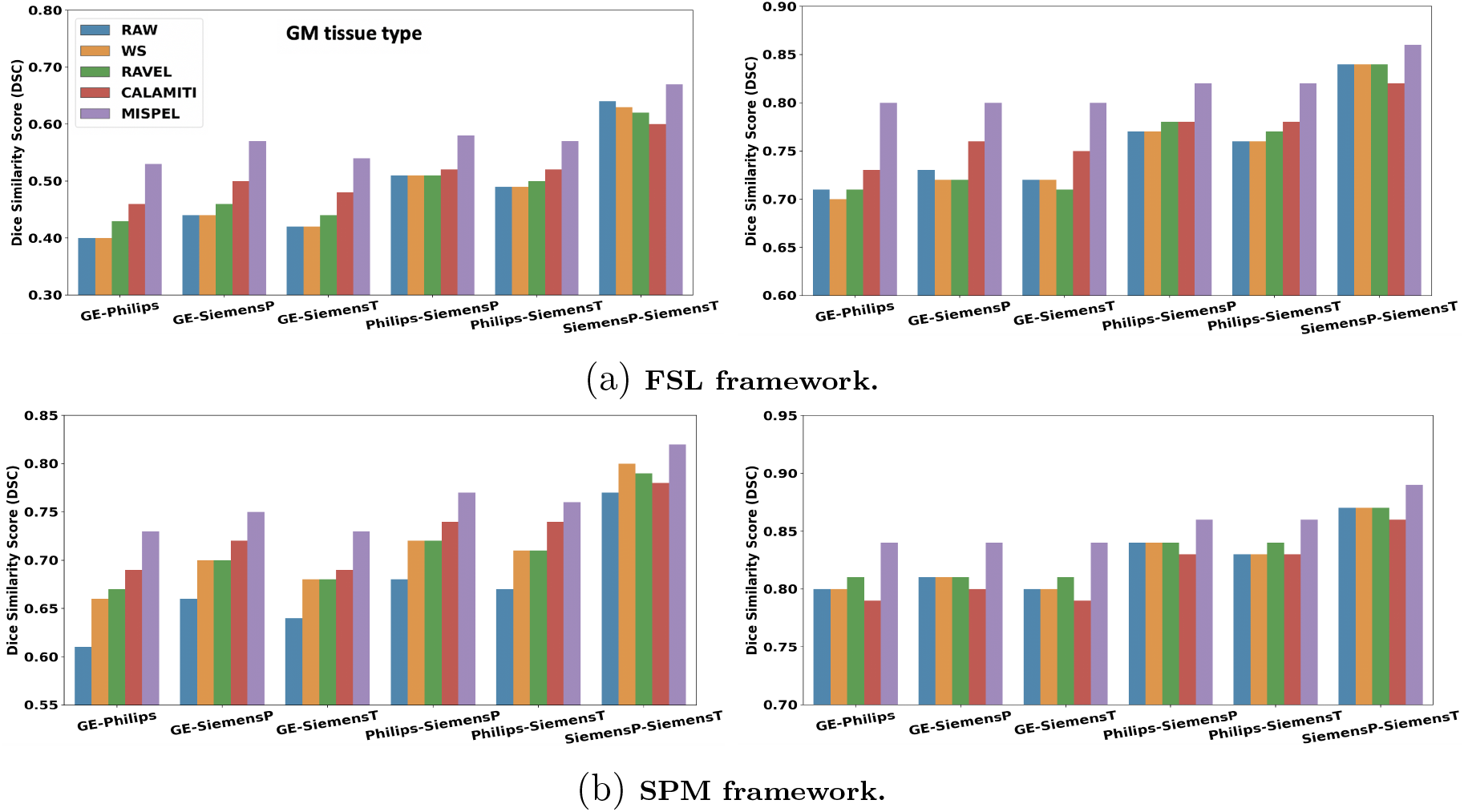
Dice similarity score (DSC) bar plots. Bar plots indicate the means of DSCs of all paired segmentations in scanner pairs. These values were calculated for segmentations of GM and WM tissue types and depicted for all four methods. These set of bar plots were depicted for volumes extracted through two segmentation frameworks: (a) FSL and (b) SPM. A harmonization method is expected to result in high mean of DSCs. MISPEL outperformed WS, RAVEL, and CALAMITI by having the largest DSC means for all scanner pairs in both FSL and SPM. For interpretation of the references to color in this figure legend, the reader is referred to the web version of this article.

We observed in Figure 10 that scanner effects exist in RAW segmentations of both FSL and SPM and appeared as relatively low means of DSC values. MISPEL outperformed WS, RAVEL, and CALAMITI in harmonization by having the largest means of DSC for all scanner pairs for both FSL and SPM. We compared the DSC distributions of MISPEL with their equivalents in RAW using paired *t* -test and all improvements of MISPEL over RAW were statistically significant. Results also showed that while WS decreased the DSC for two scanner pairs for FSL, it did better for SPM by increasing the means for 6 of the cases. RAVEL performed slightly better than WS by increasing 6 and decreasing 3 of the DSC means for FSL and improved 9 cases for SPM. CALAMITI showed 10 and 6 increases for FSL and SPM, respectively, while decreasing the rest of the cases. Using paired *t* -tests, we observed that these DSCs were statistically significantly larger than that of their RAW equivalents.

In summary, MISPEL outperformed WS and RAVEL, when FSL and SPM were used as segmentation frameworks and segmentation overlap was used as the harmonization evaluation metric.

### 3.4 Biological similarity

We investigated biological similarity of images over several biomarkers of AD: cortical thickness values of the entorhinal and inferior temporal cortices, as well as volume measures of the hippocampus and amygdala. As the evaluation criteria, we selected (1) biomarker bias and (2) biomarker variance. A harmonization method is expected to result in minimal bias and variance for the biomarkers.

#### 3.4.1 Biomarker bias and variance

Table 3 shows the biomarker bias for each of the AD biomarkers. We reported this metric for all 5 methods. For each method, we first calculated the absolute differences between paired measures of all the scanner pairs and then reported their overall mean (SD). We also compared the distribution of differences for each of the methods to that of RAW, using paired *t* -test. Moreover, Figure 11 shows the mean of RMSDs across all scanner pairs for each of the methods. These means were calculated for each of the AD biomarkers separately.

**Table 3:** Mean absolute differences for biomarkers of AD. Mean (SD) of cross-scanner absolute differences were calculated for paired measures across all scanner pairs. The measures are the FS-derived cortical thicknesses for the entorhinal and inferior temporal cortices, as well as volumes for the hippocampus and amygdala. A harmonization method is expected to decrease mean and SD of differences in RAW. MISPEL showed the best harmonization performance by having the largest number of smallest mean of differences. The distributions with the smallest means are in bold. Also, the distributions that showed statistically significant *t* -statistics when compared to RAW were marked by *.

| Mean (SD) of absolute differences over all scanner pairs |  |  |  |  |
| --- | --- | --- | --- | --- |
| Method | Cortical Thickness (mm) |  | Volume (cm<sup>3</sup>) |  |
|  | Entorhinal | Inferior Temporal | Hippocampus | Amygdala |
| RAW | 0.62 (0.42) | 0.46 (0.36) | 0.30 (0.23) | 0.25 (0.20) |
| WS | 1.00 (0.73)* | 0.63 (0.48)* | 0.43 (0.52)* | 0.23 (0.30) |
| RAVEL | 0.84 (0.57)* | 0.56 (0.41)* | 0.41 (0.29)* | 0.24 (0.21) |
| CALAMITI | 0.87 (0.60)* | 0.45 (0.32) | 0.71 (0.54)* | 0.30 (0.26) |
| MISPEL | **0.44** (0.34)* | **0.27** (0.28)* | 0.30 (0.25) | **0.17** (0.15)* |

**Figure 11:**
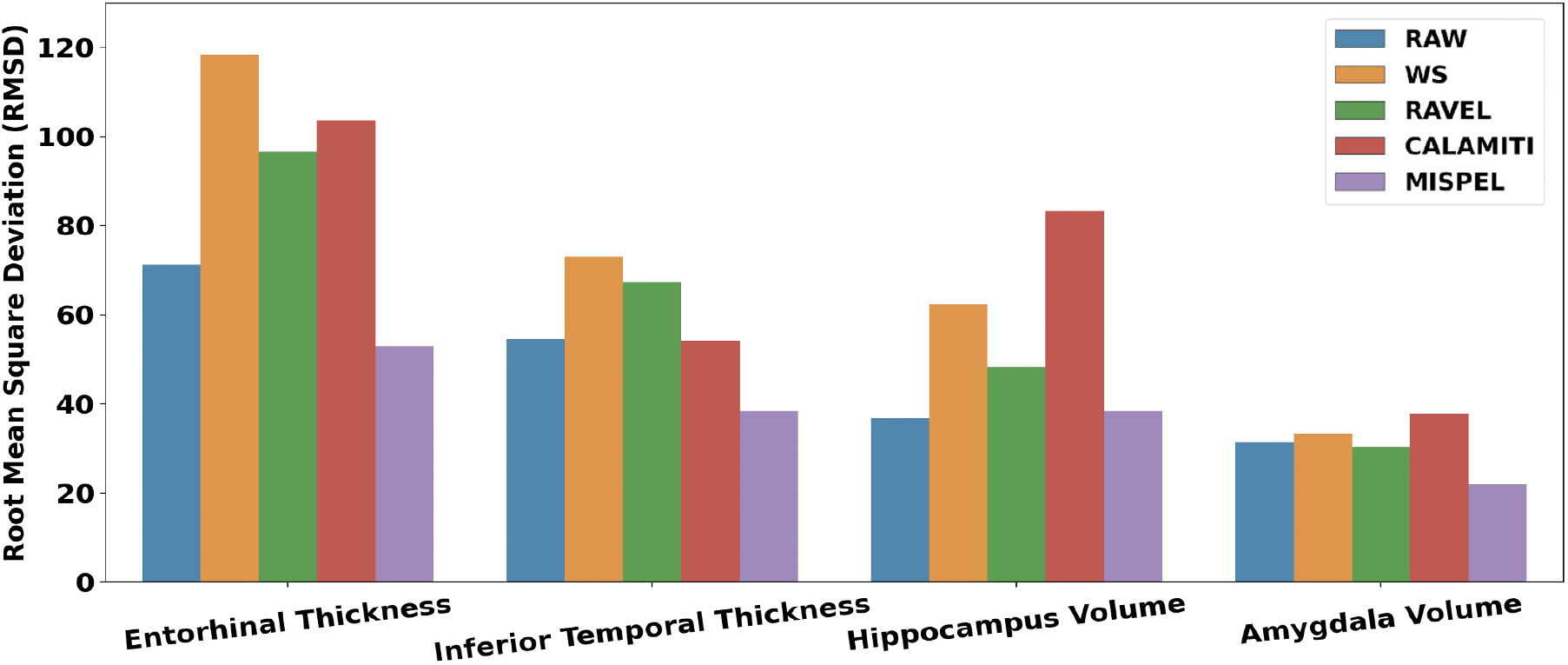
Root-mean-square deviation (RMSD) bar plots for biomarkers of AD. Each bar indicates the mean RMSD of paired measures of all scanner pairs for each of the methods. The RMSDs were reported for each of the FS-derived biomarkers of AD. A harmonization method is expected to lower values of RMSDs. MISPEL outperformed WS, RAVEL, and CALAMITI by having the largest number of smallest RMSD values. For interpretation of the references to color in this figure legend, the reader is referred to the web version of this article.

We observed in Table 3 and Figure 11 that scanner effects appeared as nonzero bias and variance values for the biomarker measures in the RAW data. We also noticed that MISPEL resulted in the largest number of statistically-significant smallest biases: 3 out of 4. MISPEL did not harmonize hippocampus, however it did not increase volumetric differences across scanners either. On the other hand, WS and RAVEL statistically significantly increased the distribution of differences for all biomarkers, except for amygdala. CALAMITI showed similar performance. This method resulted in increase in distribution of differences for 3 biomarkers while being statistically significant for 2 of them. The same trend of results was also seen for the mean of RMSD values in Figure 11.

In summary, we observed that MISPEL outperformed WS, RAVEL, and CALAMITI when harmonization was investigated as bias and variance across scanners in FS-derived biomarkers of AD.

### 3.5 Analysis on biological variables of interest

We investigated whether harmonization could succeed in preserving or strengthening SVD-related group differences in our data. For this, we studied the Cohen’s d effect sizes of SVD groups in each of the scanners. We calculated these values for each of the biomarkers and methods separately. Table 4 shows mean (SD) of these Cohen’s d values over all scanners. A harmonization method is expected to not reduce these means of Cohen’s d after harmonization, that is to preserve group differences. We observed that MISPEL increased effect sizes for all of the biomarkers, except for hippocampus. MISPEL resulted in a minor decrease in Cohen’s d of hippocampus. On the other hand, WS, RAVEL, and CALAMITI resulted in major decreases for hippocampus and amygdala, a minor decrease for inferior temporal, and a minor increase for entorhinal. In summary, we observed that MISPEL succeeded in preserving our biological signal of interest and outperformed other methods in this respect.

**Table 4:**
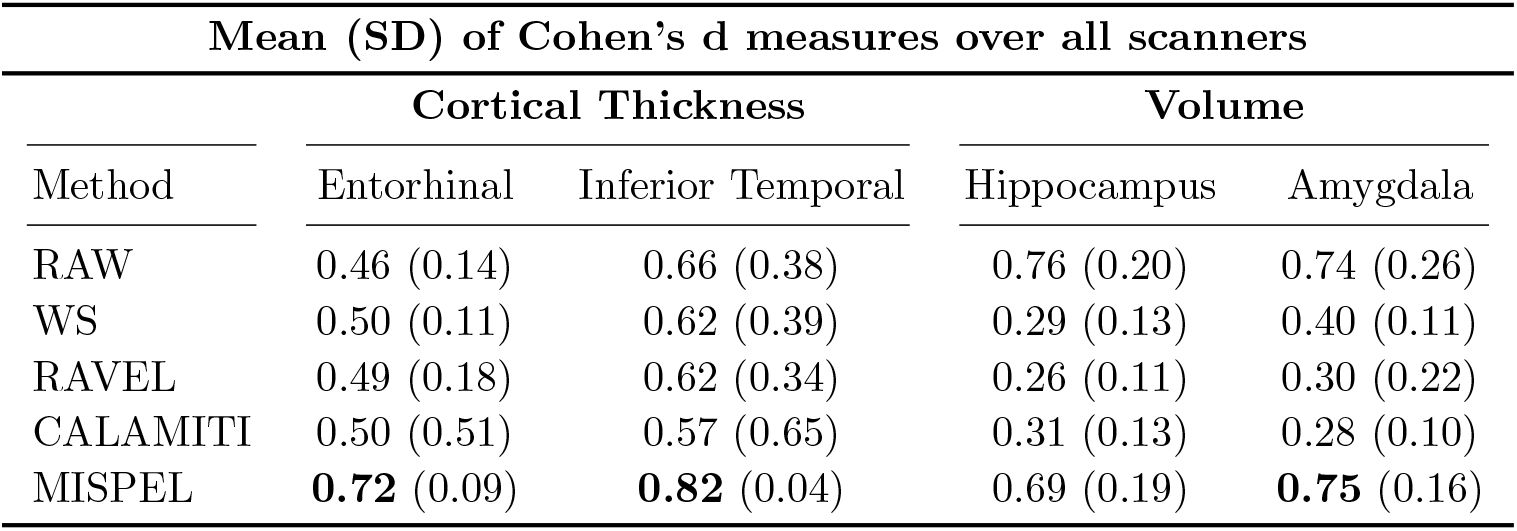
Mean (SD) of Cohen’s d measures for bimarkers of AD. Mean (SD) of Cohen’s d values were calculated over all scanners for biomarkers of AD and all methods. A harmonization method is expected to preserve or increase the effect sizes calculated relative to RAW. Increased effect sizes relative to RAW are in bold.

## 4 Discussion

In this study, we presented MISPEL, a supervised deep harmonization technique for removing scanner effects from images of multiple scanners, while preserving their biological and anatomical information. Unlike other supervised or unsupervised methods, MISPEL is a multi-scanner method mapping images to a scanner *middle-ground* space in which images are harmonized. We evaluated MISPEL against commonly used intensity normalization and harmonization methods (White Stripe, RAVEL, and CALAMITI) using a set of evaluation criteria including image similarity, GM-WM tissue contrast, tissue volumes and segmentation similarity, and biological similarity in a dataset of matched T1 MR images acquired from 4 different 3T scanners. We also investigated whether these methods could preserve or even enhance the SVD group differences as a biological signal of interest. We found that (1) scanner effects appear in our dataset as dissimilarity in image appearance/contrast, GM-WM contrast, tissue type volumetric and segmentation distributions, and distributions of regional measures of AD; (2) White Stripe normalized images, but did not achieve harmonization; (3) RAVEL and CALAMITI achieved harmonization to some extent; and (4) MISPEL outperformed all other methods in harmonization.

Based on the evaluated harmonization metrics, we observed that images of GE were more similar to those of Philips and images of SiemensP showed more similarity to SiemensT’s. We also observed that scanner effects appeared mainly as the dissimilarity between pairs of GE or Philips and SiemensP or SiemensT. We observed that removing intensity unit effects using White Stripe successfully normalized images (Supplementary Figure 1) and resulted in improved image similarity, but did not majorly enhance other metrics we used for evaluating harmonization. The relative failure to harmonize may be due to the fact that WS is an intensity normalization method, which does not account for scanner information. We also observed that WS increased the variability of image-derived measures across subjects. Such behavior was observed in bias and variance metrics for GM and WM volumes, as well as biomarkers of AD. This was expected as WS is an individual-level method. This property of WS makes the normalization of any new unseen image more convenient but may also result in inconsistent normalization across images. WS also decreased the effect size for volumetric biomarkers of AD, when SVD group differences were studied. In fact, scaling and centering the intensity distributions does not necessarily remove scanner effects; on the contrary, over-matching distributions could result in the removal of other sources of variability that could be of interest (Fortin et al., 2016). These results show that scanner effects are not addressed solely through intensity normalization and a more comprehensive harmonization method is necessary.

RAVEL is an unsupervised normalization and harmonization framework that could extract components of scanner effects for each of the subjects as inter-subject variability across their CSF area. Our results show that RAVEL achieved harmonization to some extent relative to White Stripe, but was outperformed by MISPEL. RAVEL increased the similarity of images in their appearance/contrast, GM-WM contrast, and tissue type volumes and segmentation overlap when the SPM framework was used. However, RAVEL could not achieve harmonization for FSL-derived GM and WM volumes. Moreover, it deteriorated the bias and variance for biomarkers of AD, except for volumes of the Amygdala. RAVEL also did not preserve the SVD group differences when *volumetric* biomarkers were investigated. These relative failures could be due to several reasons. First, RAVEL uses neither the information of scanners nor the matched data during its harmonization process. Second, RAVEL is prone to remove some biological variability across subjects, if such variability is not accounted for in RAVEL modeling. RAVEL also showed large variability and inconsistent harmonization across subjects, especially for FSL-derived volumes. Such results have been also reported in (Torbati et al., 2021a) when RAVEL was used for harmonizing paired images of GE 1.5T and Siemens 3T scanners and FreeSurfer was used. Similar results were observed for WS. Thus, such behavior of RAVEL could be due to using WS in its normalization step.

For a fair comparison with CALAMITI, we used it in a supervised manner by applying it to our inter-scanner paired dataset instead of inter-modality paired data as discussed in Zuo et al. (2021). Results showed that CALAMITI achieved harmonization to some extent relative to White Stripe. However, it did not perform better than RAVEL and was outperformed by MISPEL. CALAMITI improved similarity of images in their appearance/contrast, GM-WM contrast, and tissue type volumes and segmentation overlap when the SPM framework was used. CALAMITI did not show consistent harmonization for FSL-derived volumes. It resulted in both increased and decreased biases for these measures. Moreover, CALAMITI showed large variability and inconsistent harmonization across subjects for both FSL- and SPM-derived volumes. This method did not achieve harmonization for AD biomarkers either. It deteriorated the bias and variance for the entorhinal and hippocampus measures. It also deteriorated the SVD group differences for all biomarkers, except for the entorhinal. These failures in harmonization could be due to CALAMITI’s harmonization approach. CALAMITI encodes paired images into their mutual scanner-invariant anatomical components, and their individual contrast and scanner-variant components. For harmonizing an image, it synthesizes the harmonized image by using its anatomical component and the target scanner’s contrast component/scanner component. Such methodology is prone to losing some anatomical information of images, if it could not segregate the anatomical and contrast components properly. Similar harmonization failures were observed for CALAMITI in (Zuo et al., 2021) when image-derived summary measures were investigated.

MISPEL outperformed White Stripe, RAVEL, and CALAMITI based on all harmonization evaluation criteria. MISPEL mapped images to a middle-ground harmonized space, in which matched images were made more similar in contrast by removing scanner effects. For our data, GE and Philips images were more similar to those of SiemensP and SiemensT, in terms of GM-WM contrast and tissue type volumetric distributions. It should be noted that no directed mapping or a *target* scanner was selected for MISPEL harmonization, and MISPEL does not require a selected *target*. In fact, MISPEL naturally finds this middle-ground space. GE and Philips images were made more similar to SiemensP and SiemensT, with relatively minimal change made to SiemensP and SiemensT by MISPEL, likely due to SiemensP and SiemensT images being most similar and therefore biasing the middle-ground space found by MISPEL. For this scenario of data, not requiring a target scanner could be regarded as an advantage for MISPEL over other deep-learning based harmonization frameworks. Other widely used statistical harmonization methods, including WS, RAVEL, and ComBat, also do not require a target scanner. However, harmonizing to a middle-ground rather than a specified target could be problematic in other scenarios, such as if the data were collected on a majority of lower-quality scanners. This may bias MISPEL to learn a lower-quality middle-ground space for harmonizing images and degrade the quality of images from more advanced scanners. In such cases, MISPEL could potentially be modified to map images to a target scanner.

Results from volumetric and segmentation evaluations also show that MISPEL image-based harmonization improves the harmonization of downstream image analysis results regardless of framework. It showed improvement for both segmentation platforms tested, FSL and SPM, which have been shown to largely differ in their segmentation results (Tudorascu et al., 2016) even in healthy volunteers. MISPEL also showed success in harmonization of biomarkers of AD and enhancing the SVD group differences when these biomarkers were used. The improved performance of MISPEL compared to RAVEL and CALAMITI could be due to the design choices for MISPEL. First, U-Net (Ronneberger et al., 2015) units were used as the encoder-decoder units in MISPEL. The U-Net could preserve the structure of brain by transferring the information of images from encoder layers to the decoder layers. Second, the loss functions for MISPEL were selected cautiously to tackle the contrast discrepancy within paired images and preserve their anatomy. Even so, MISPEL is far from perfect. We observed that MISPEL showed better harmonization for cortical thickness biomarkers relative to volumetric measures. MISPEL improved volumetric bias and variance for the amygdala and preserved the SVD group differences in amygdala volumes, but MISPEL also reduced the SVD group differences in hippocampal volumes.

One possible reason for the suboptimal performance of MISPEL in hippocampal-derived harmonization metrics could be related to its 2D network. Such a network may result in slice-to-slice inconsistency for harmonized images. To evaluate this, we assessed slice-to-slice *consistency* measures for each of the RAW and MISPEL-harmonized images. We collected an array of SSIM measures between each adjacent axial slice of each image. We then paired each of the harmonized images with their equivalent RAW image and calculated the correlation between SSIM consistency measures of images of each pair. A harmonization method that preserves the slice-to-slice consistency of RAW images should have a statistically significant correlation near 1 over all pairs. We conducted this experiment for slices of each brain orientation separately and observed 0.994 (ranges: [0.969, 0.999]), 0.992 (ranges: [0.962, 0.999]), and 0.992 (ranges: [0.973, 0.999]) mean of correlations across subjects for axial, sagittal, and coronal slices, respectively. These high correlations demonstrate that slice-to-slice inconsistency is not a significant concern for MISPEL when trained exclusively on axial slices. As such, further investigation is necessary to optimize MISPEL for multi-scanner studies where focal regional volumes are of interest.

Our study adds to the growing harmonization literature by (1) presenting MISPEL, a supervised multi-scanner harmonization method; (2) introducing a multiscanner matched dataset of four 3T scanners; (3) providing a set of experiments assessing scanner effects and evaluating harmonization; and (4) evaluating the practical harmonization performance of MISPEL against widely used and state-of-the-art image intensity normalization and harmonization methods. One limitation of our study is the use of a single matched-scan cohort. The generalizability of MISPEL to unmatched multi-scanner data, relative to existing and commonly used normalization and harmonization methods, was not assessed. As future work, we will investigate whether MISPEL harmonization can be improved for volumetric measures by using the 3D fusion network proposed in (Zuo et al., 2021). This network synthesizes the harmonized image by fusing the 2D harmonized slices of the image across all orientations. We will also study the generalizability of MISPEL to other matched datasets with different degrees of scanner effects, such as paired GE 1.5T and Siemens 3T data (Torbati et al., 2021a), as well as unmatched multi-scanner datasets. We will further study MISPEL across other modalities, such as Fluid-attenuated inversion recovery (FLAIR). Using these new datasets and modalities, we will investigate whether not selecting a target scanner for MISPEL could result in suboptimal harmonization and whether modifying MISPEL to map images to the space of a target scanner could improve image quality.

In this article, we proposed a supervised multi-scanner harmonization approach, MISPEL, that harmonizes the T1-w MRI of scanners for which a matched dataset is available. The main design goal for MISPEL was preserving the anatomical information of images while harmonizing them. MISPEL showed decent harmonization performance while our well-suited set of evaluation criteria was used. This set uses the matched data to investigate harmonization from various aspects. MISPEL and our evaluation criteria are promising tools to help multi-site studies dealing with the scanner technical variability.

## Credit authorship contribution statement

Mahbaneh Eshaghzadeh Torbati: Conceptualization, Methodology, Writing – original draft. Davneet S. Minhas: Methodology, Writing – review & editing. Charles M. Laymon: Writing – review & editing. Pauline Maillard: Writing – review & editing. James D. Wilson: Writing – review & editing. Chang-Le Chen: Writing – review & editing. Ciprian M. Crainiceanu: Methodology, Writing – review & editing. Charles S. DeCarli: Funding acquisition, Writing – review & editing. Seong Jae Hwang: Funding acquisition, Methodology, Conceptualization, Supervision, Writing – review & editing. Dana L. Tudorascu: Funding acquisition, Methodology, Conceptualization, Supervision, Writing – review & editing.

## Acknowledgments

This work was supported by the following NIH/NIA grants: R01 AG063752 (D.L. Tudorascu), P30 AG10129 and UH3 NS100608 (C. DeCarli), and the University of Pittsburgh Alzheimer’s Disease Research Center Grant P30 AG066468 (S. Hwang). S. Hwang was also supported by Institute of Information & communications Technology Planning & Evaluation (IITP) grant funded by the Korea government (MSIT), Artificial Intelligence Graduate Program, Yonsei University, under Grant 2020-0-01361-003, and the Yonsei University Research Fund of 2022 (2022-22-0131).

## 5 Appendix

**Supplementary Figure 1:**
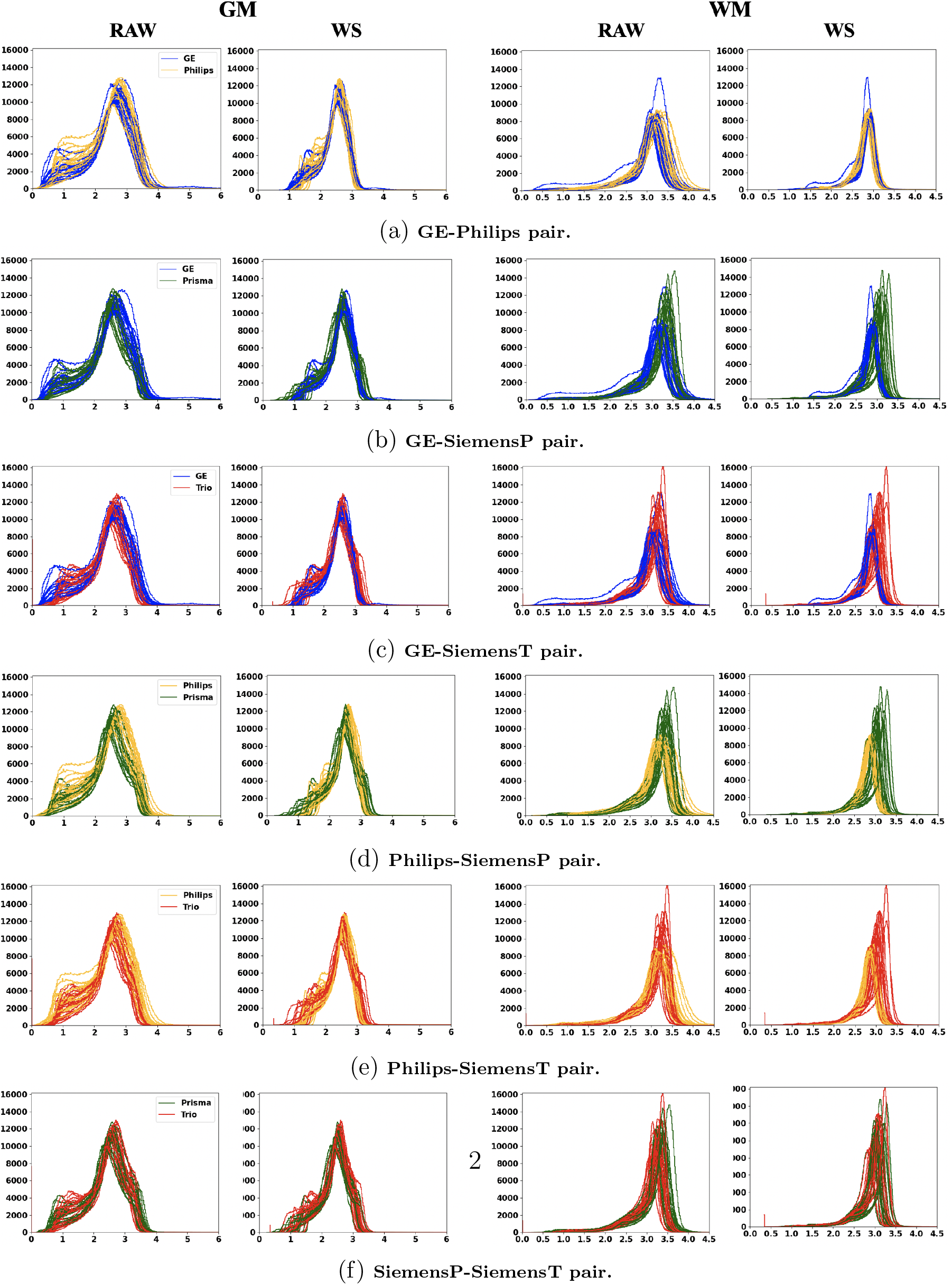
Histograms of gray matter (GM) and white matter (WM) voxels for RAW and White Stripe (WS)-normalized images of all subjects. These histograms were plotted for all 6 scanner pairs. WS makes the plots more centered, overlapped and therefore comparable across subjects. WS usually outputs images with negative intensity values. for plotting the histograms, we shifted the WS-normalized images to have positive intensity values. For interpretation of the references to color in this figure legend, the reader is referred to the web version of this article.

## Footnotes

1 https://github.com/Mahbaneh/MISPEL

